# Human prostate cancer bone metastases have an actionable immunosuppressive microenvironment

**DOI:** 10.1101/2020.03.19.998658

**Authors:** Ninib Baryawno, Youmna Kfoury, Nicolas Severe, Shenglin Mei, Karin Gustafsson, Taghreed Hirz, Thomas Brouse, Elizabeth W. Scadden, Anna A. Igolkina, Bryan D. Choi, Nikolas Barkas, John H. Shin, Philip J. Saylor, David T. Scadden, David B. Sykes, Peter V. Kharchenko, as part of the Boston Bone Metastasis Consortium

## Abstract

Bone metastases are devastating complications of cancer. They are particularly common in prostate cancer, represent incurable disease and are refractory to immunotherapy. We sought to define distinct features of the bone marrow microenvironment by analyzing single cells from prostate cancer patients’ involved bone, uninvolved bone and distant bone sites as well as bone from cancer-free, orthopedic patients and healthy individuals. Metastatic prostate cancer was associated with multifaceted immune distortion, specifically exhaustion of distinct T cell subsets, appearance of macrophages with states specific to prostate cancer bone metastases. The chemokine CCL20 was notably overexpressed by myeloid cells, as was its cognate CCR6 receptor on T cells. Disruption of the CCL20-CCR6 axis in mice with syngeneic prostate bone metastases restored T cell reactivity and significantly prolonged animal survival. Comparative high resolution analysis of prostate cancer bone metastasis shows a targeted approach for relieving local immunosuppression for therapeutic effect.

## Introduction

Prostate cancer frequently metastasizes to bone where, in the castration-resistant setting, 70 to 90% of patients have radiographically-detectable skeletal involvement (de Bono et al., 2011; de Bono et al., 2010; Halabi et al., 2016; Petrylak et al., 2004; Tannock et al., 2004). To date, bone metastases represent a generally incurable form of prostate cancer and contribute significantly to disease-specific morbidity and mortality (Xiang and Gilkes, 2019). In particular, metastatic involvement of the vertebral bodies can result in pathologic fracture and spinal cord compression leading to significant neurologic and functional disability, often necessitating urgent or emergent surgical decompression and stabilization. Surgical decompression/stabilization and targeted radiation are used selectively. Approved bone-targeted systemic therapies offer some benefit: zoledronic acid and denosumab delay skeletal morbidity but do not improve overall survival (Saad et al., 2004). The radiopharmaceutical radium-223 offers a modest improvement in overall survival (Parker et al., 2013).

Spinal bone marrow has long been known to be a primarily hematopoietic organ and represents a unique reservoir for several types of immune cells such as macrophages, dendritic cells (DCs), myeloid derived suppressor cells (MDSCs), and varied T cell subsets that has the ability to dramatically influence the trajectory of malignant disease. The critical role of the innate myeloid immunity in the tumor microenvironment goes well beyond its classical role of phagocytosis and antigen presentation. In addition to shaping the tumor adaptive immune response, myeloid cells impact response to cancer therapy (De Palma and Lewis, 2013; Engblom et al., 2016; Mantovani et al., 2017), promote angiogenesis (De Palma et al., 2017; Lewis et al., 2016a) and directly contribute to tumor progression and metastasis through the secretion of growth factor and extracellular matrix degrading enzymes (Coffelt et al., 2016; Kitamura et al., 2015; Lewis et al., 2016a). Despite this, major roadblocks such as our incomplete understanding of myeloid cellular heterogeneity and plasticity in addition to major differences between mouse and human myelopoiesis to name few, hinder the efficient translation of these findings towards improved disease outcomes.

The immune checkpoint therapies targeting CTLA4 and PD-1/PD-L1 that have proven to be very effective against a wide range of tumors, have thus far have been unsuccessful in clinical trials in unselected prostate cancer populations (Laccetti and Subudhi, 2017) despite the high abundance of T lymphocytes in the tissue. In particular, emerging evidence indicates that patients with bone metastases are at a disadvantage when it comes to response to immune checkpoint therapies (Beer et al., 2017). This may in part be explained by the unique composition of the bone marrow T cell population. Immunosuppressive T regulatory cells (Tregs) are particularly abundant in the marrow space (Zou et al., 2004) and display a more activated phenotype (Glatman Zaretsky et al., 2017). In addition, when naïve T helper cells (T_H_) become activated in the bone marrow, the local cytokine environment appears to favor the development of T_H_17 over T_H_1 responses – a process that has detrimental impact on the success of immune checkpoint therapy (Jiao et al., 2019). Altogether, this indicates that there is a clear need for new and more effective approaches for our patients with bone metastatic prostate cancer based on a better understanding of the immune microenvironment that is permissive of its growth and is a critical player in response to therapy.

Though the bone marrow microenvironment is clearly hospitable to prostate cancer, there have been multiple longitudinal barriers to gaining a better understanding of the supportive relationships between cell types. Bone biopsies present challenges centered on patient discomfort, feasibility, and sample quality after decalcification. Imaging biomarkers of bone metastasis response to systemic therapy are substantially limited. Preclinical models of prostate cancer metastatic to bone have historically been limited as well. Finally, bulk sequencing of tumor tissue has yielded important insights but is limited by its inability to characterize sub-populations and specific expression of ligands and receptors of tumor, immune, and stromal cells. In current clinical practice, the tumor tissue that is removed surgically in cases of spinal cord compression has little diagnostic utility. In this study, we now systematically collect this fresh tissue for cell isolation, phenotyping, and expression analysis with droplet-based single-cell RNA sequencing to facilitate analysis of rare populations of immune, tumor, and stromal cells in this environment. We hypothesize that an improved understanding of immune cell support of malignant cells within the bone marrow will identify areas of vulnerability amenable to therapeutic intervention.

## Results

### Widespread alteration of the bone marrow by prostate cancer metastasis

Availability of fresh clinical prostate cancer bone metastatic samples is essential to the study of relationships between cancer cells and the marrow microenvironment. Bone sampling by needle biopsy can be technically challenging and provides scant quantities of tissue. We therefore developed an inter-disciplinary workflow to obtain tissue from emergent clinically-indicated surgeries for those rare cases of spinal cord compression due to epidural extension of tumor or pathologic fracture of a vertebral body. The collection of tissue samples was integrated into the standard workflow of surgery. We collected matched sets of tissue fractions from each patient: solid metastatic tissue (*Tumor* fraction), liquid bone marrow at the vertebral level of spinal cord compression (*Involved*), as well as liquid bone marrow from a different vertebral body distant from the tumor site but within the surgical field (*Distal*) (Fig. 1A, Supplementary Table 1). This allowed for a comparison within the same individual, controlling for inter-individual variation. All patients had a historic diagnosis of prostate cancer and had standard pathologic evaluation to confirm prostate cancer in the bone marrow within tissue sampled at the time of spinal decompression surgery (Fig. 1B, S1A). Additionally, each had a preoperative magnetic resonance imaging (MRI) of the spine showing obvious evidence of tumor causing spinal cord compression (Fig. 1C, S1B). Bone marrow samples from patients undergoing hip replacement surgery (*Benign*) served as a non-malignant comparator group. The transcriptional composition of all samples was assessed using single-cell RNA-seq (scRNA-seq). The measurements yielded on average of 2,490 cells per sample, detecting on average 3952 molecules per cell (Supplementary Table 2). The 10x Chromium V2 protocol utilized captures a relatively modest percentage of mRNA molecules in each cell (Ding et al., 2020; Mereu, 2020; Wang, 2020). However, its ability to measure thousands of individual cells in each sample enables one to distinguish cell subpopulations and more subtle differences associated with changes in cellular state through downstream computational analysis (Lambrechts et al., 2018; Peng et al., 2019; Zhang et al., 2019).

**Figure 1.**
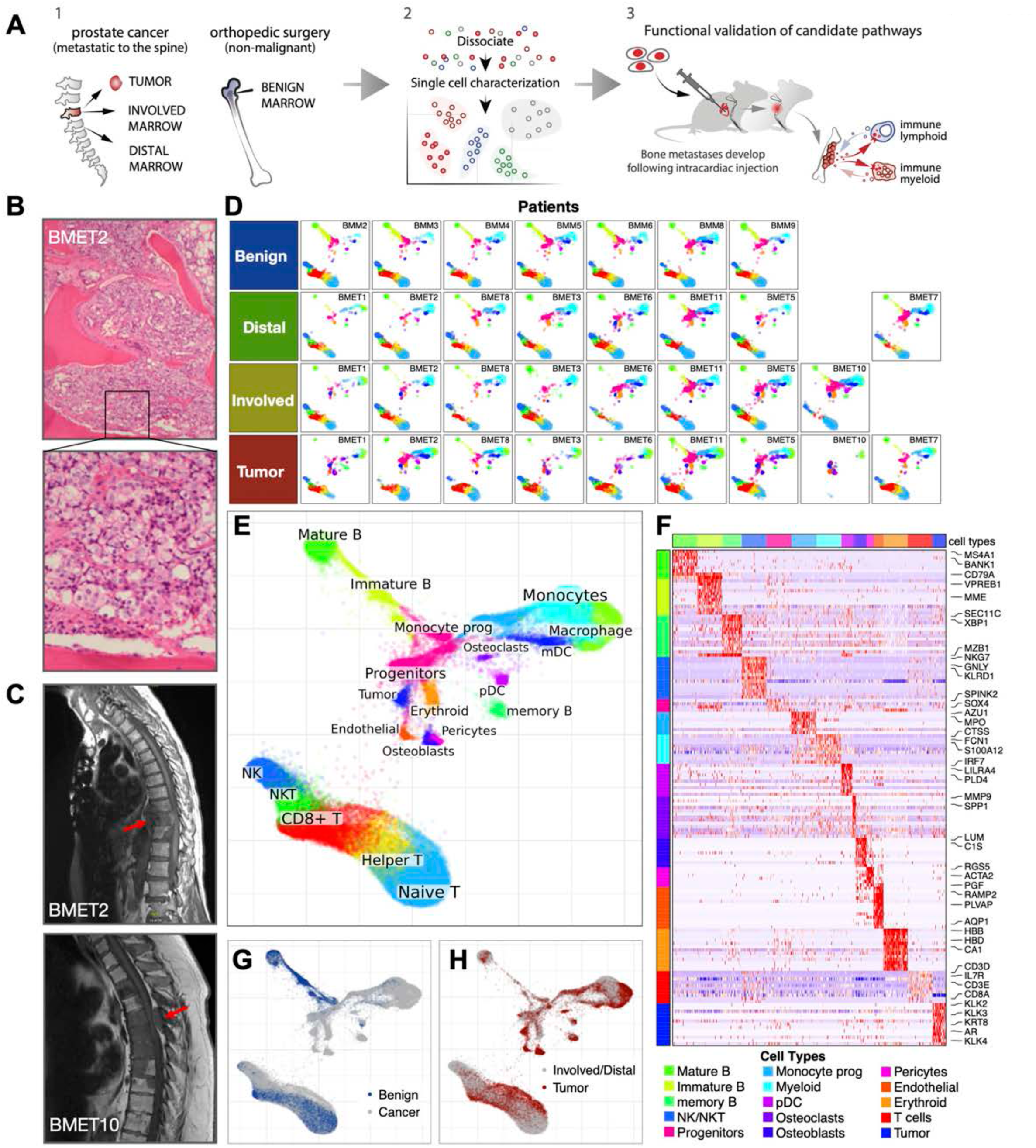
Transcriptional impact of prostate cancer metastasis on the bone marrow. **A**. Schematic illustration of study design. **B**. H&E-stained tissue sections of metastatic tumor resected from a representative patient with bone metastatic prostate cancer (100x -top, 400x -bottom). **C**. Sagittal T1 MRI imaging of the thoracic spine for each of those three cases showing tumor masses with spinal cord compression. (top) The T10 vertebral body (arrow) is structurally compromised by tumor with pathologic fracture and retropulsion of bone fragments into the ventral epidural space. *(bottom)* An extradural tumor mass in the dorsal spinal canal at the TS level (arrow) with diffuse tumor involvement of vertebral bodies of TS - T7. **D**. The analysis integrates scRNA-seq from three fractions (Tumor, Involved, Distal) of 9 metastatic prostate cancer patients (rows), as well as 7 Benign bone marrow controls from hip replacement surgery patients. The datasets were integrated to establish joint embedding and annotation (E). Projections of individual samples are shown in D. **F**. Marker genes for major cell populations. **G**. The difference in frequency of major subpopulations between Benign controls and cancer patients (all three fractions). **H**. Compositional differences between Tumor and Involved/Distal fractions.

We used a recently-developed approach to integrate the full collection of samples (Fig. 1D,E) (Barkas et al., 2019). Joint analysis revealed a rich repertoire of immune cells and ongoing hematopoiesis with HSC/progenitor populations giving rise to B cell, monocyte, and erythroid lineages (Fig. 1F, Supplementary Table 3). The T cell population was disconnected (Fig. 1E), which is expected as T cell maturation occurs outside of the bone marrow within the thymus. Granulocytes, whose abundance was quantified using flow cytometric analysis (Fig. S1D), were absent in the single-cell data, reflecting a known cell lineage limitation of the single-cell protocol (Chen et al., 2018). The non-immune subpopulations, mostly captured within the *Tumor* fraction, included endothelial cells, pericytes, osteoblasts, osteoclasts as well as the tumor cells themselves (Fig. 1E).

The tumor cells, which were annotated based on expression of key prostate cancer markers (KLK4, KLK2, AR) (Fig. 1F), displayed patient-specific chromosome-scale deviations of expression magnitudes indicative of the presence of CNVs (Patel et al., 2014; Tirosh et al., 2016) (Fig. S2A), and notable inter-individual variation of expression patterns. Tumor cells were also detected in some of the *Distal* fractions, reflecting diffuse marrow involvement in certain patients (Fig. S2B,C). Tumor cells exhibited strong patient-specific expression differences (Fig. 2A), however, analysis of intra-tumoral heterogeneity revealed four major aspects of tumor cell variation that were shared by different patients (Fig. 2B-D, S2D-F). Three of the aspects (IC2-4) reflected variation in the metabolic activity related to protein, nucleic acid, and ribosomal metabolism. The remaining aspect (IC1), however, was driven by transcription of genes associated with cell differentiation and AP-1 signaling, including IER2, JUNB, and SOX4, that have all been shown to facilitate motility and invasion of metastatic cells (Bilir et al., 2016; Hyakusoku et al., 2016; Neeb et al., 2012; Tiwari et al., 2013). This aspect was strongly enriched for functions related to vasculature development, epithelial cell differentiation, as well as regulation of hematopoietic regulation (Fig. 2E). This AP-1 related aspect of intratumoral variation distinguished clearly separated subpopulations within the metastatic tumors of multiple patients (Fig. 2F).

**Figure 2.**
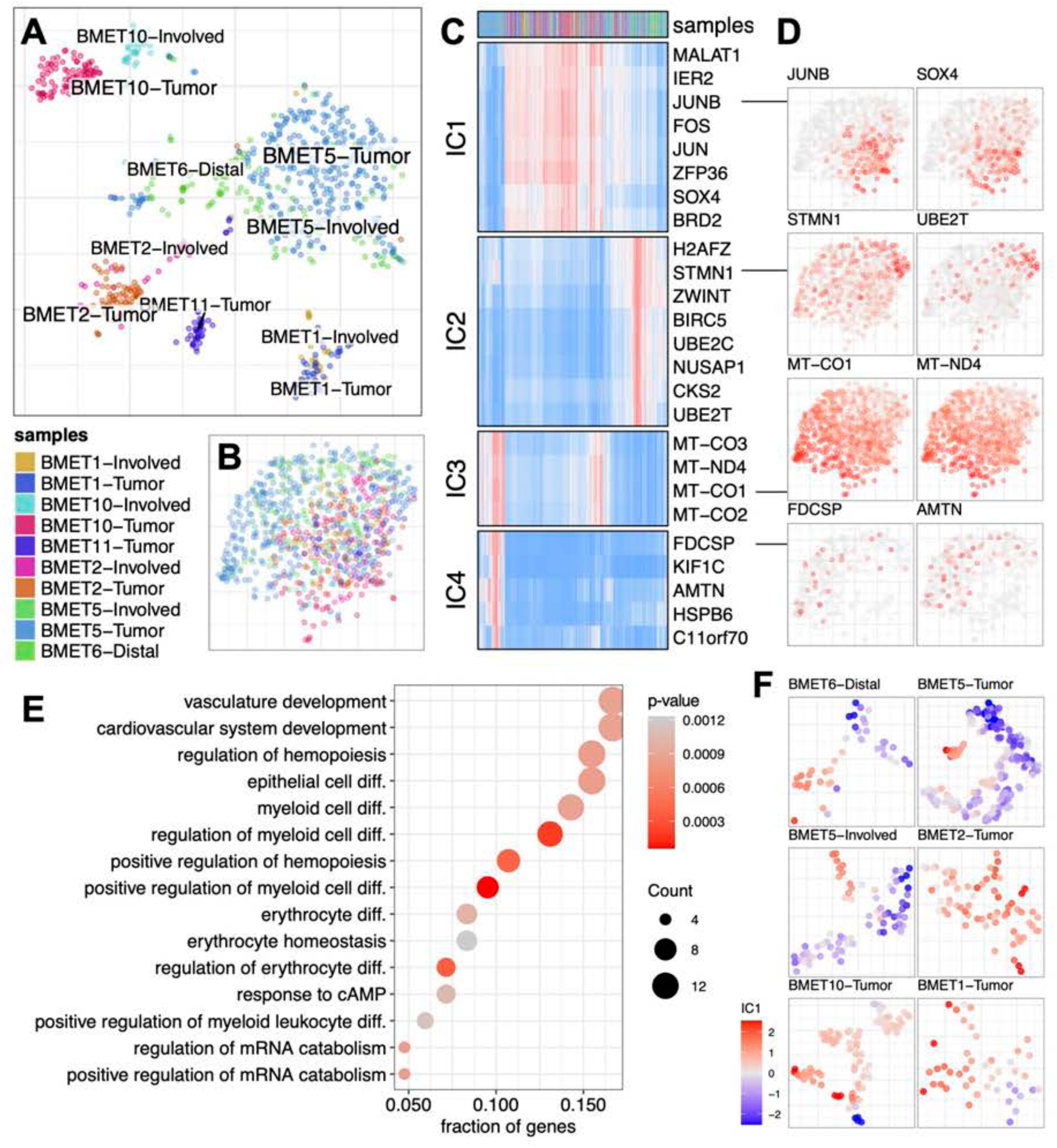
Common axes of intratumoral heterogeneity are seen within the metastatic tumor populations. **A**. Transcriptional heterogeneity of tumor cells from different samples is visualized on a tSNE embedding, showing strong patient-specific patterns. **B**. Integration of tumor cells from different patients using Conos (same colors as in **A). C**. Expression of high-loading genes for four independent components (ICs) is shown based on the ICA analysis of the integrated tumor cell populations, illustrating re-current aspects of transcriptional heterogeneity between patients (red -high expression; blue -low). **D**. Expression of high-loading genes identified by ICA are shown on the combined Conos embedding (B) (red -high expression; white -low). **E**. Gene Ontology terms enriched in the top 200 high-loading genes of the IC1 aspect of transcriptional heterogeneity. **F**. The IC1 cell scores are shown for a subset of samples using sample-specific t-SNE embeddings (without integration). The IC1 component separates major cell subpopulations visible within the individual samples.

The presence of metastases significantly altered the immune cell composition of the bone marrow. Relative to *Benign* controls, the three cancer fractions showed a striking depletion of B cells and B cell progenitors (Fig. 1G, 3B, S1D, S3A). The *Tumor* fraction had an increase in the proportion of macrophages compared to the *Involved/Distal* fractions (Fig. 1H). As expected, the solid *Tumor* fraction also showed more abundant endothelial, pericyte, osteoblast, and tumor cell populations (Fig. 1H, S3A). As analysis of simple cell proportions can be skewed by a single large-scale change, such as depletion of a large B cell compartment, we confirmed the shifts mentioned above using a more robust compositional data analysis technique (see Methods, Fig. S3D).

Complementary to the shifts in the frequency of cell types, scRNA-seq data also captured the transcriptional state permitting an examination of expression differences (see Methods). With the exception of the *Benign* controls, the overall similarity of cell states in different samples reflected patient-specific signatures (Fig. S3B). The extent of inter-patient differences was smallest for the *Benign* controls, and incrementally increased towards the site of metastasis (*Distal* to *Involved* to *Tumor* fraction) (Fig. 3C). This significant increase in the inter-patient variability in cancer, and the *Tumor* fraction in particular, demonstrates the divergent impact of metastasis on the bone marrow across individuals. Nevertheless, we find that for most cell types the expression difference between cancer and control groups significantly exceeds the magnitude of inter-individual variation within the groups (Fig. S3C), underscoring the marked impact of the presence of the metastasis on the transcriptional states of different populations. The HSC/progenitor population (*Progenitors*) was among the most affected. Analysis of differentially expressed genes within this heterogeneous population revealed pronounced downregulation of cell cycle in the metastatic context, complemented by upregulation of translation and immune activation functions (Fig. 3D,E, S3E,F). While the specific signatures of the state differences varied, this general pattern of functional impact was mirrored by most other cell types, with many cell types also exhibiting upregulation of stress response pathways in the metastatic context (Fig. 3D,E). Observing such broad impact of metastasis across different cellular compartments, we focused our analysis on the detailed changes affect the two major immune compartments, myeloid and T-cells. For this purpose, we excluded two patients (BMET-10 and BMET-11) that had received Radium 223 treatment (Supplementary Table 1), as the detailed patterns of expression signatures appeared to be distinct. The general trends described above, however, remained consistent with or without the inclusion of these two patients.

**Figure 3.**
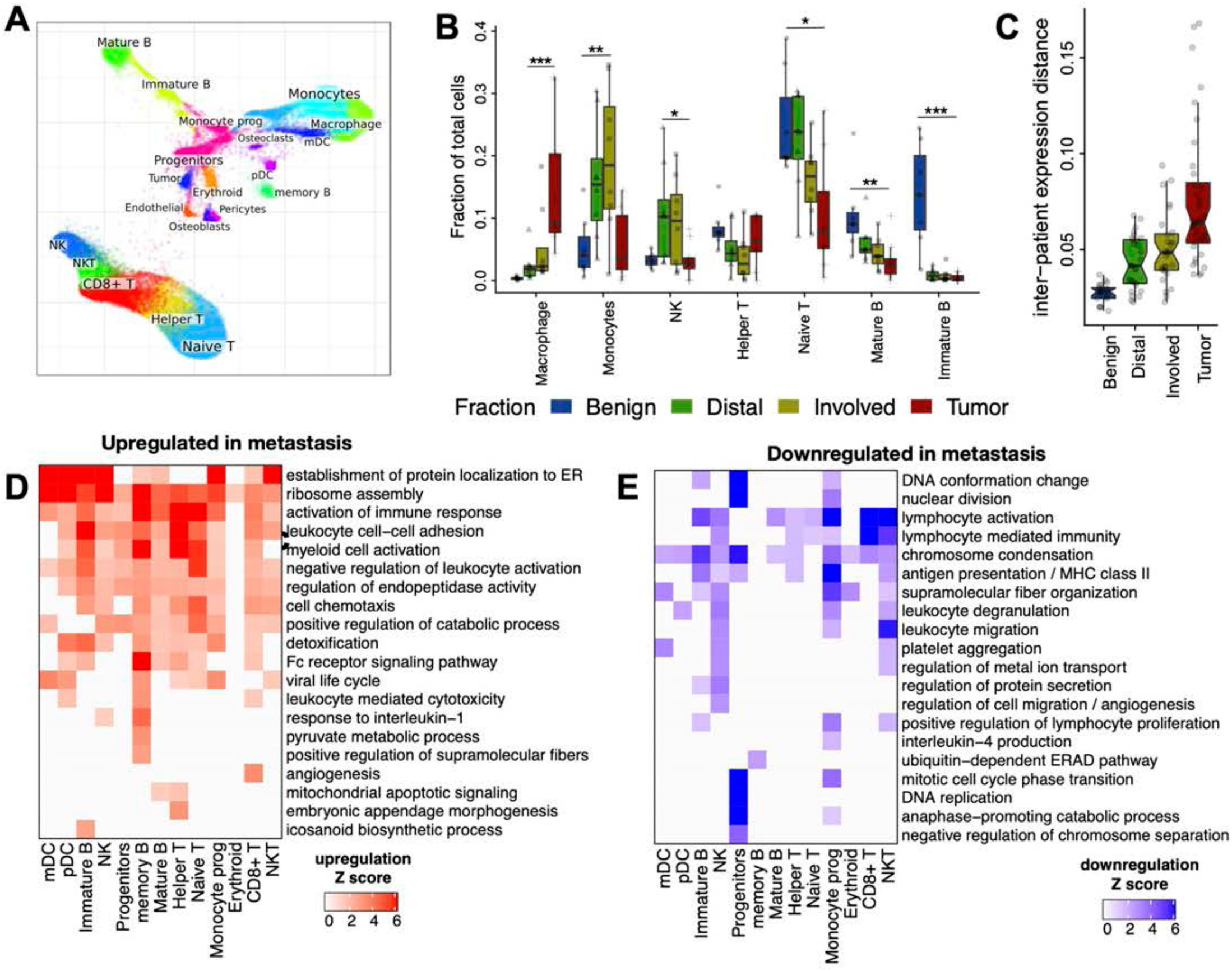
Overview of compositional and transcriptional shifts associated with presence of metastasis in the bone marrow. **A**. Visualization of major cell populations from all sample fractions on a joint embedding (same as Fig. 1E). **B**. Compositional shifts for the most affected cell types are shown as boxplots comparing proportion of major cell populations between Benign controls and different cancer fractions (see Fig. S3A for all populations). Statistical significance is shown for select pairs, based on a Wilcoxon rank sum test, reported at 99% reproducibiilty power based on bootstrap resampling of cells (*p<0.05, ****p<0.0001). **C**. Average cell-type-specific expression distances between different patients is shown for Benign controls and the three cancer fractions. The analysis shows that controlling for the compositional differences, the expression state of different cell types in Benign controls is significantly more similar than in the cancer fractions. The expression differences between individuals increase incrementally towards the site of metastasis, with the highest expression variation observed in the Tumor fraction. **D**,**E**. Overview of the GO BP category enrichment in the top 300 up-regulated (D) and down-regulated (E) genes in the Involved *vs*. Benign differential expression analysis performed for each major cell subpopulation. Gene sets showing significant enrichment (FDR <0.05) were clustered into 20 clusters according to the similarity of the participating genes, and for each cluster (row) the name of the GO category with the lowest FDR is shown.

### Inflammatory monocytes and immunosuppressive macrophages in tumors

The myeloid cells have been implicated in supporting tumor progression in certain cancers (Coffelt et al., 2016; Lewis et al., 2016b). Focused examination of the myeloid compartment revealed substantial population shifts between *Benign* and malignant fractions (Fig. 4A,B, S4A,B). While *Benign* controls primarily contained resting monocytes, the monocytes of both *Involved* and *Distal* fractions expressed genes indicative of an activated and proliferative state in cancer patients (Mono-2, myeloid cell activation, GO:0002275 P<10^−16^). Striking changes were observed in the *Tumor* compartment, driven by the appearance of prominent populations of Tumor Inflammatory Monocytes (TIMs), and Tumor Associated Macrophages (TAMs) (Fig. 4B-D, S4B,C, Supplementary Table 4), a finding that was validated by flow cytometric analysis of independent samples (Fig. 4E, S1D, Fig. S4D).

**Figure 4.**
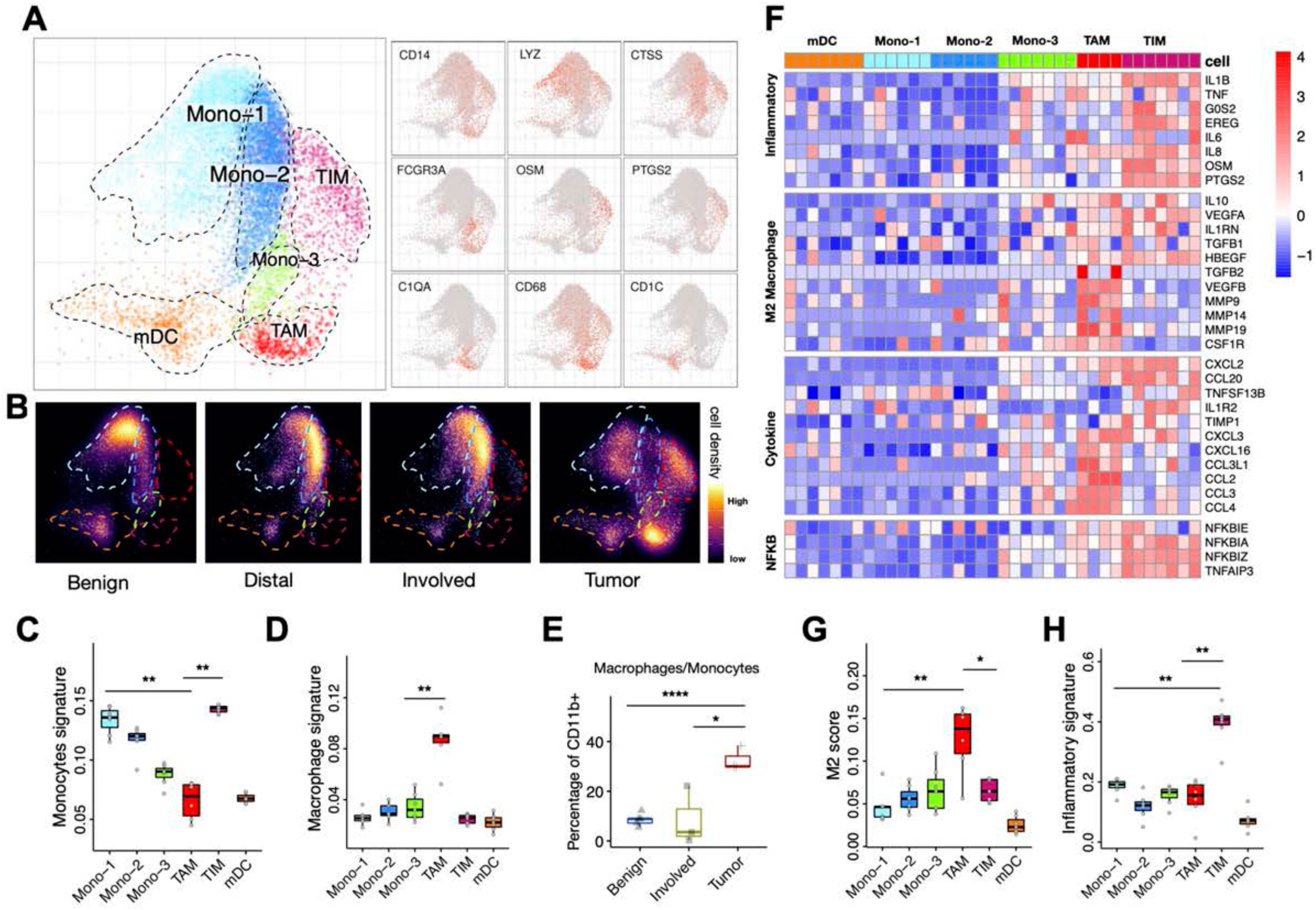
Myeloid populations shift towards inflammatory monocytes and repressive macrophages in the Tumor fraction. **A**. Detailed annotation of the myeloid subpopulations is shown on a myeloid-specific joint embedding {left), together with select gene markers (right). **B**. Changes in the composition of the myeloid compartment between sample fractions is visualized as cell density on the joint embedding. The composition of the Distal and Involved fractions is dominated by an activated Mono-2 population, while Tumor fraction shows appearance of Tumor Inflammatory Monocyte (TIM) and Tumor Associated Macrophages (TAM) populations. **C**,**D**. Average expression of monocyte (C) and macrophage (D) signaturegenes (see Methods, GSE5099) in different myeloid subpopulations. Statistical significance was assessed using Wilcoxon test, reported at 99% reproducibility power (see Methods). **E**. Boxplot showing the percent of monocytes/macrophages within CD11b+ cells (Supplementary Table 7) by flow cytometry in Benign, Involved and Tumor fractions from three independent patients. Statistical analysis was performed using Student’s t-test (*p<0.05, ****p<0.0001). **F**. Heatmap shows average expression of select genes from different categories (rows) across different cell populations (top color bar, colors matching panel A) for each patient (columns). **G**,**H**. Average expression of the M2 macrophage signature gene panel (G), and monocyte inflammatory gene panel (H) across different monocyte populations.

The TAM population had an expression pattern characteristic of M2 macrophages (Azizi et al., 2018) (Fig. 4F,G) which have been shown to suppress anti-tumor immune responses across a broad range of tumors (DeNardo and Ruffell, 2019). Accordingly, TAM cells express anti-inflammatory cytokines (TGFB1, IL-10, Interleukin 1 receptor antagonist IL1rn) (Sanjabi et al., 2009) as well as factors implicated in cancer growth and invasiveness (CCL2, CCL3, CCL4, VEGF, MMPs) (Bachelder et al., 2002; Merchant et al., 2017; Pellikainen et al., 2004) (Fig. 4F). Interestingly, TGFB1 has been recently shown to be a molecular mediator for T helper cell polarization toward Th17 in response to immune checkpoint therapy in the bone metastatic microenvironment and blocking it in combination with the immune checkpoint inhibitors improved survival in a mouse model of prostate cancer (Jiao et al., 2019). Further, a recent study by an independent group suggest that exposure to the anti-androgen drug enzaluatamide is associated with an increased tumor-intrinsic activation of TGFβ signaling and EMT signature in metastatic castration resistant prostate cancer (He, 2020), suggesting that resistance to anti-androgen treatment may be induced by TGFB1. In contrast to the TAM immunosuppressive M2-like signature, TIM cells had a pro-inflammatory monocyte signature with high expression of pro-inflammatory cytokines such as IL1-B and TNF (Fig. 4F,H, S4G) (Becking et al., 2015; Smillie et al., 2019).

The TIM and TAM populations that we observed in primary samples closely paralleled *ex vivo* effects on peripheral blood monocytes of tumor-secreted factors (Vlaicu et al., 2013). The cells expressed epidermal growth factor-like ligands (epiregulin (EREG) in TIMs, HB-EGF in TAMs) and a common repertoire of interleukin-6-like JAK/STAT3 pathway activators (IL-6, Oncostatin-M (OSM)) (Fig. 4F). Both signaling modalities support tumor growth directly as well as indirectly by suppressing the activation of CD8+ T cells, expanding suppressive Treg cells, promoting M2 macrophage polarization and differentiating myeloid cells into myeloid derived suppressor cells (MDSCs) (Huynh et al., 2017). TIMs and TAMs also expressed inhibitors of the pro-inflammatory nuclear factor-*k*B pathway (NFKBIA, NFKBIZ, TNFAIP3) (Lawrence, 2009) (Fig. 4F). The presence of these TIM and TAM expression signatures are associated with significantly worse prognosis for patients across a broad range of cancer types (Fig. S4E).

Comparison with published single-cell RNA-seq data shows that cells analogous to TIM and TAM subpopulations can also be found in other cancers (Lambrechts et al., 2018; Peng et al., 2019; Zhang et al., 2019), with the highest abundance of TIM and TAM-like cells observed in liver and lung cancer, respectively (Fig. 5A,B). Their molecular state, however, differs significantly from that observed in our data. The TIM population we have identified in the prostate-origin metastasis shows significantly stronger inflammatory signature, compared to other examined cancers (Fig. 5C,E). For TAM, the M2 polarization observed in the metastatic context was comparable to those seen in other tumors (Fig. 5D), however the expression of most cytokines mentioned above differed in the bone metastatic context profiled here (Fig. 5F). Similarly, both TIM and TAM populations showed prominent nuclear factor-*k*B signaling signature atypical of the other cancers. As there is currently little data on other metastatic cases, especially in the bone, studies of other cancers will be needed to determine to what extent the observed alterations of the bone marrow are specific to metastasis of prostate origin.

**Figure 5.**
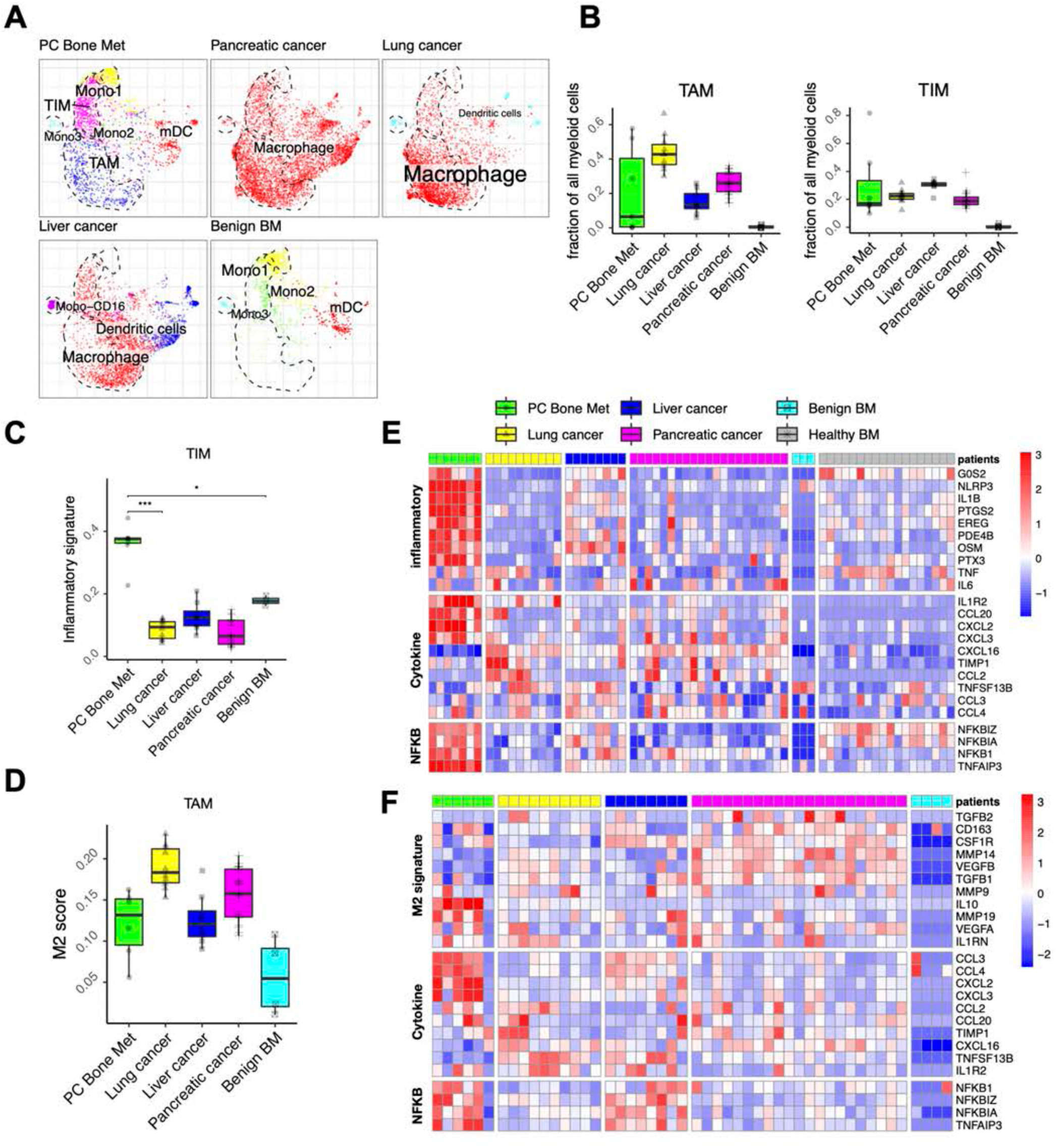
Comparative of myeloid states with other cancers. **A**. Joint alignment of myeloid cells from benign bone marrow, prostate cancer bone metastasis, pancreatic cancer, lung cancer and liver cancer, colored by original cell annotation. The top left box shows the cells collected from the prostate metastatic patients (this study), with contours outlining the major meyloid cell populations. The same contours are shown for comparison in all other panels, which show from other cancers (and Benign control from this study in the last panel). **B**. Boxplot showing relative abundance of TIM and TAM across different cancer types. **C**,**D**. Average expression of monocytes inflammatory signatures (C) and M2 macrophage signatures (D) in different cancer types. **E**,**F**. Heatmap shows average expression of select genes from different categories (rows) across different cancer types (top color bar, colors matching panel A) for each patient (columns), respectively for TIM (E) and TAM (F).

### Expansion of dysfunctional cytotoxic T lymphocyte subpopulation in bone metastases

Detailed analysis of the lymphocyte compartment revealed the expected subsets, CD4^+^ T helper (T_H_) and T regulatory (Treg) cells, CD8^+^ cytotoxic T lymphocytes (CTLs), as well as NK and NKT cells (Fig. 6A, Fig. S6A, Supplementary Table 3, 4). The smallest population of CD8^+^ T cells expressed high levels of *CCR7, LEF1* and *SELL*, indicating that these are antigen-inexperienced, naïve cells (Fergusson et al., 2016; Picker et al., 1993; Willinger et al., 2006) (Fig. S6A). The CTLs could be further subdivided into two subtypes: CTL-1 expressing effector T cell genes such as *KLRG1, GZMK* and other cytotoxicity mediators, whereas CTL-2 are characterized by an effector/memory-like transcriptional profile including *IL7R* and *KLRB1* (Fig. 6B) (Fergusson et al., 2016; Zhang et al., 2018). We also identified CCR7 expressing naïve CD4+ cells, whereas the mature T helper cell compartment appeared to be a mix of T_H_1 and T_H_17 cells (T_H_1/17) expressing *CXCR3* and *CCR6* respectively (Fig. S6A) (Zhang et al., 2018). Unfortunately, the data did not provide sufficient resolution to further distinguish these two populations.

**Figure 6.**
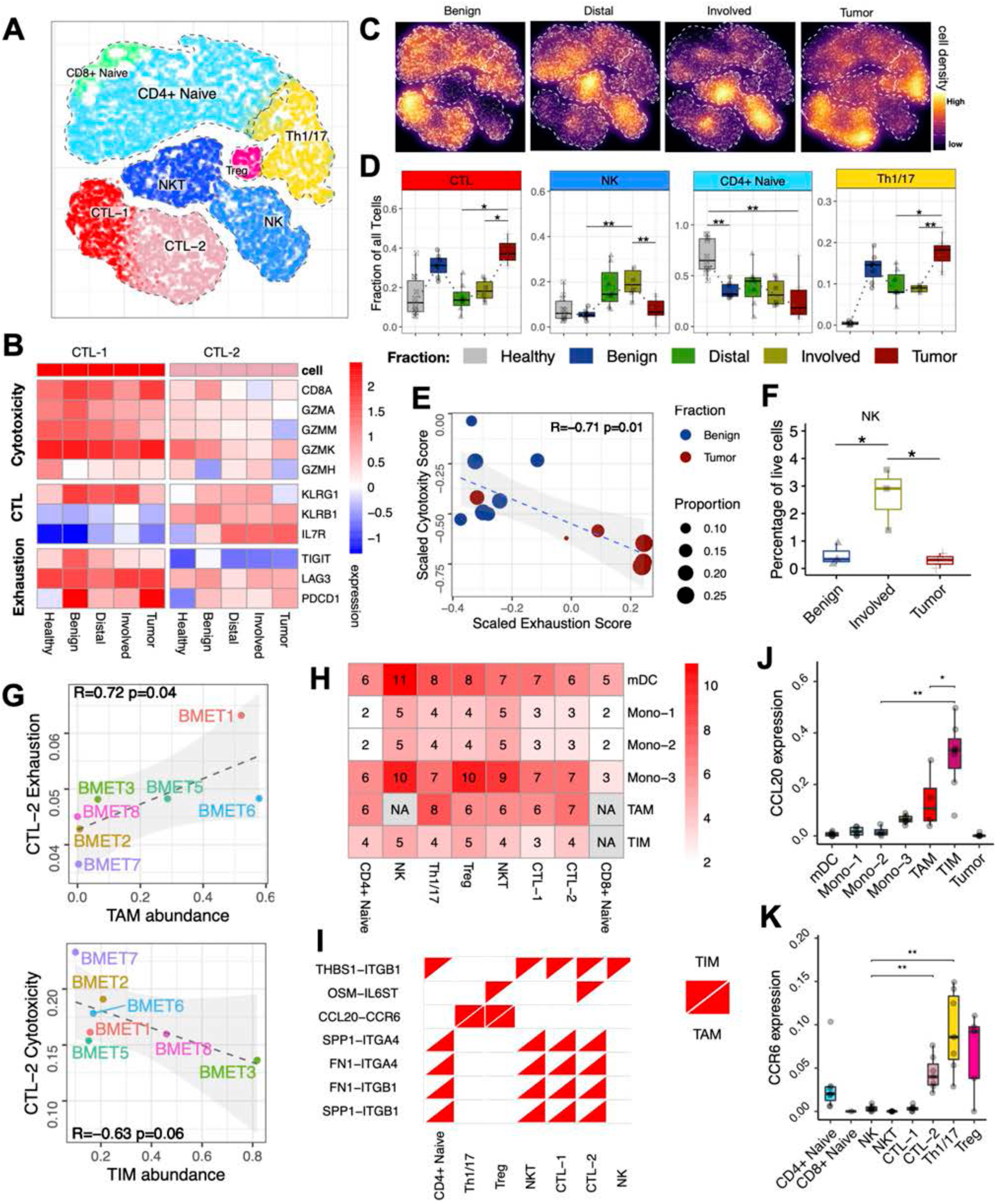
Tumor fraction shows increased abundance of exhausted cytotoxic populations and helper T cells. **A**. Detailed annotation of the T cell subpopulations, shown on a T-cell specific joint embedding. **B**. Expression of key genes from different categories (rows) in CTL-1 and CTL-2 populations is shown across different sample types. **C**. Shifts in the T cell populations across Benign and cancer fractions is visualized as cell density on the joint embedding. **D**. Boxplots of cell frequency changes of different subpopulations between sample fractions (including Healthy controls, grey). **E**. Cytotoxicity and exhaustion scores for CTL-1 in Benign and Tumor fraction (see Supplementary Table 4). Points size shows proportion of the CTL-2 cells relative to all T cells in the sample. Statistical significance was assessed using Wilcoxon test, reported at 99% reproducibility power. **F**. Flow cytometry validation of the NK cell abundance (Supplementary Table 7) in Benign, Involved, and Tumor fractions from three independent patients. Statistical analysis was performed using a Student’s t-test (*p<0.05). **G**. The abundance (x axis) ofTAM (lower) and TIM (upper) populations s associated with the increased exhaustion and decreased cytotoxicity of the CTL-2 populations (y axis). Pearson linear correlation estimate and p-values are shown. **H**. The number of known cognate receptor-ligand pairs for which the ligand is expressed in the myeloid subpopulation (rows), and the ligand is expressed in the T cell population (columns). NAs represent pairs of populations that were observed in fewer than 5 patients. **I**. Receptor-ligand channels connecting TIM and TAM populations to the T cells, based on additional filtering criteria (see Methods). **J**,**K**. Cluster-average expression of CCL20 (J) and CCR6 (K) is shown for different cell population.sStatistical significance was assessed using Wilcoxon test, reported at 99% reproducibility power.

The relative abundance of the CTL, and T_H_ / T_H1/17_ populations followed a common pattern, gradually increasing from the *Distal* fraction towards the site of metastasis (*Tumor* fraction) (Fig. 6C,D), whereas other T-cell types were found to be unchanged (*e*.*g*. Tregs) or decreased in the Tumor (*e*.*g*. naïve T-cells) (Fig 6C,D, S6B). The abundance of activated effector T cells and Tregs was also high in the *Benign* controls (Fig 6C,D, S6B), likely reflecting the known inflammation in the osteoarthritis patients from whom the samples were collected. An additional set of bone marrow samples from *healthy* individuals (Oetjen et al., 2018) revealed significantly lower abundance of these populations, offset by higher proportion of CD4^+^ naïve cells (Fig. 6D). The functional state of these T cell populations was also impacted, in particular CTL-2 in the *Tumor* fraction exhibited reduced cytotoxicity expression signature in combination with a pronounced T cell exhaustion signature, characteristic of dysfunctional CTL commonly observed in tumors (Lee et al., 1999; Li et al., 2019) (Fig. 6E, S6C-E,I). The level of T cell exhaustion observed in the CTL-2 cells of the *Tumor* fraction was significantly higher than that of the *Benign* controls, highlighting the distinction between sustained benign inflammation and the metastatic context. The NK cell population showed a complementary pattern, with high abundance in the *Involved* and *Distal* samples, but not in *Benign* and *Tumor* fractions (Fig. 6C,D) - an observation that was further validated by flow cytometric analysis (Fig. 6F, S1E). This suggests that although NK cells can be recruited to the general area of the bone metastasis, they may fail to infiltrate the tumor.

### Coordination between myeloid and lymphoid compartments

The observed deficiencies of cytotoxicity in the proximity of the tumor may arise through repressive actions by other immune populations (Munn and Bronte, 2016). For instance, we find that the Treg cells, which typically act to suppress immune response (Woo et al., 2002), showed increased activity signatures at the site of the metastasis (Fig. S6F-H). The complex patterns of immune signaling are also likely to involve the myeloid compartment. Indeed, considering variation of TAM and TIM abundance among patients (Fig. S4F) we find that increased proportion of TAMs at the site of metastasis is correlated with CTL-2 exhaustion (Fig. 6G). Similarly, increased abundance of TIMs coincides with a reduction in the CTL-2 cytotoxicity signature (Fig. 6G). While these associations suggest that TIM or TAM populations may be directly affecting T lymphocyte state, identifying a specific signaling channel through which communication takes place is challenging.

There are currently no effective computational or high-throughput experimental methods for carrying out such inference. The space of potential interaction channels is extensive: screening a database of ligand and cognate receptors (Efremova et al., 2020) for those expressed in myeloid and T lymphoid compartments, respectively, we find 241 potential channels (39 for TAM, 29 for TIM, Fig. 6H, Supplementary Table 5). To prioritize likely candidates, we applied additional filtering criteria, requiring up-regulation of the receptor expression in *Tumor* in comparison to *Benign* controls, and high levels of ligand expression in the corresponding myeloid population, reducing the number of potential channels to 7 (Fig. 6I). Of the cytokines that were expressed within the TIM and TAM cells of the *Tumor* fraction, CCL20 was one of the most highly expressed (Fig. 4F, 6J, S4G), and its cognate receptor CCR6 was predominantly expressed on Treg and T_H_1/17 cells (Fig. 6K, S7A). Studies of other cancers have demonstrated that CCL20 signaling can promote tumor growth, invasiveness and chemoresistance (Lee et al., 2017; Lu et al., 2017; Walch-Ruckheim et al., 2015) by recruitment of Tregs and/or T_H_17 cells (Walch-Ruckheim et al., 2015; Wang et al., 2019). CCL20 expression on the protein level was confirmed with ELISA using plasma from different bone marrow fractions (Supplementary Table 6). CCL20 has been shown to be expressed on primary prostate cancer tumor cells and in stromal fibroblasts within the tumor microenvironment (Beider et al., 2009; Walch-Ruckheim et al., 2015). However, in the metastatic samples, CCL20 was expressed primarily by the TIM and TAM populations, and was absent from the tumor cells (Fig. 6J). Overall, the direct role of the CCL20-CCR6 axis in human prostate bone metastases is unclear.

### CCL20-CCR6 signaling leads to T lymphocyte exhaustion

To investigate the impact of CCL20-CCR6 signaling axis in prostate bone metastasis, we developed a syngeneic model of prostate bone metastasis, based on the RM1-BM cell line derived from C57BL/6 RM1 prostate cancer cells (Power et al., 2009; Thompson et al., 1989). These cells were sequentially injected into C57BL/6J wild-type mice to generate a highly penetrant bone tropic prostate cancer cell line, RM1-BoM3, that induced osteolytic and osteoblastic lesions as shown by micro-computed tomography (microCT) and radiography (Fig. S7B-E). Using this model, we demonstrate that the absence of the CCR6 receptor improved survival of bone metastases bearing mice when compared to controls (Fig. 7A,B). Likewise, administration of CCL20 blocking antibody to wild-type mice with prostate cancer bone metastasis resulted in a survival advantage in comparison with isotype treated controls (Fig. 7C). Single-cell analysis on T cells from the bone metastasis isolated from wild-type and CCR6-KO mice showed average reduction in Active Tregs and Exhausted CTLs with a concomitant increase in Naive CTLs (Fig. 7D, S7F).

**Figure 7.**
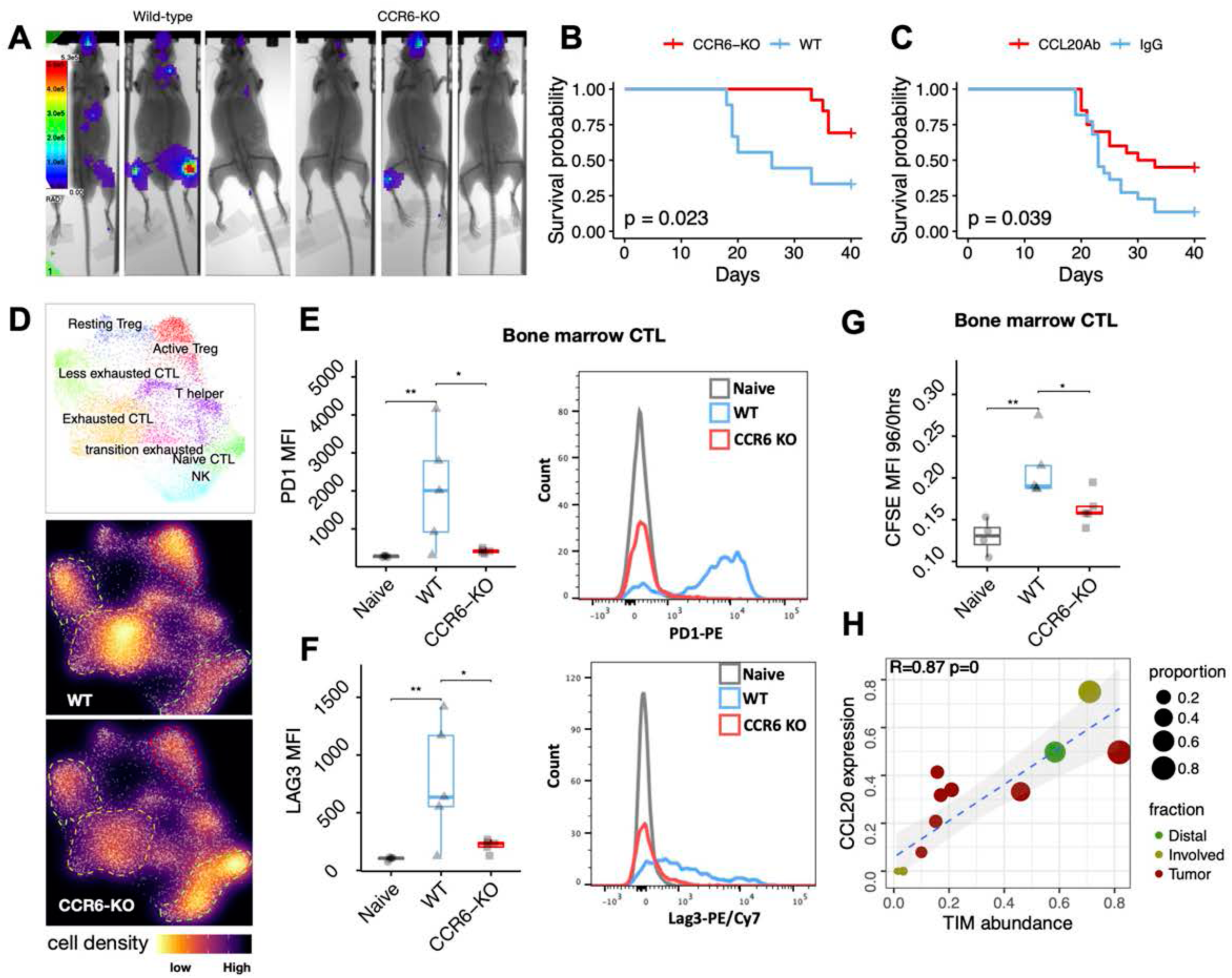
Disruption of the CCL20-CCR6 signaling axis relieves T cell exhaustion and improves survival. **A**. Representative images of bone metastases localization as shown by bioluminescence optical imaging in the mandible or in long bones of syngeneic wild-type and CCR6-KO mice one week post injection of RM1-BoM3 cells. **B**. CCR6 knockout results in a statistically significant survival benefit in syngeneic mouse model of bone marrow metastasis (n=9 wild-type and n=13 CCR6-KO). Two independent experiments (Supplementary Table 8). **C**. Block of CCL20 by an antibody results in a statistically significant survival benefit in syngeneic mouse model of bone marrow metastasis (n=22 wild-type and n=20 CCL20Ab). Two independent experiments (Supplementary Table 8). **D**. CD3+ T cells were isolated from wild-type and CCR6-KO mice and analyzed by single-cell RNA-seq (n=2 mice per group). Changes in the composition of the T cell compartment between wild-type and CCR6-KO is visualized as cell density on the joint embedding. **E**,**F**. Flow cytometric analysis of T cell exhaustion markers PD-1 (F) and Lag-3 (H) on CTLs from naïive, and tumor bearing wild-type and CCR6-KO mice. (n= 5 mice per group). **G**. Dilution of CFSE in sorted CD8+ CTLs was analyzed by flow cytometry after 96 hours of stimulation and compared to baseline levels as an indicationof T cell proliferation. (n= 4 naïve, n= 5 wild-type, n= 5 CCR6-KO) **IH** The magnitude of CCL20 expression in the TIM population (y axis) is strongly correlated with the overall abundance of TIM cells, measured as a proportion of all myeloid cells (x axis).

These results suggest that blockade of the CCR6-CCL20 axis counteracts immunosuppression in the bone metastasis tumor microenvironment. In order to further confirm this impact on the CTLs, flow cytometric analysis of T cell populations was performed on bone marrow from bone metastasis bearing wild-type and CCR6-KO mice as well as tumor naive wild-type mice. This revealed a significant increase in the frequencies of both CD4+ T_H_ cells and CD8+ CTLs in the marrow of CCR6-KO mice with prostate cancer bone metastasis (Fig. S7I). The infiltrating CD8+ CTLs also appeared to be less exhausted in the CCR6-KO mice as evidenced by reduced cell surface levels of exhaustion markers PD-1 and Lag3 (Fig. 7E,F). Notably, the CCR6-KO CTLs were found to display a similar exhaustion marker profile as their tumor naive counterparts, suggesting that CTL activity is unimpeded in the tumor microenvironment after inhibition of CCR6-CCL20 signaling (Fig. 7E,F). To test this, CD8+ CTLs were sorted from wild-type and CCR6-KO mice with tumor as well as from tumor naive wild-type mice and stimulated *in vitro*. After 96 hours of stimulation, wild-type CTLs proliferated at a significantly lower rate than both tumor naive and CCR6-KO cells (Fig. 7G, S7J), indicating that increased expression of exhaustion markers displayed by these cells, translates into a reduced functionality as well.

## Discussion

Together, these results provide a high-resolution landscape of the human bone marrow in the context of metastatic prostate cancer and show that it is distinctive from either the immune activated state of individuals requiring hip replacement for osteoarthritis or of healthy controls. The cancer affected state has a broad but now defined set of signature changes in the hematopoietic cells of the bone marrow. These include a decrease in the cell cycle status of primitive hematopoietic cells and a decrease in B cells and B cell progenitors. The similarity of the *Involved* and *Distal* fractions suggests that these differences likely reflect systemic changes accompanying cancer. In contrast, increases in monocyte/macrophage and the CTL and T_H1/17_ T cell subpopulations were more evident in the sites of tumor involvement. These provided evidence for TIM and TAM populations expressing activated signatures and candidate molecules that might affect the T cell populations and the exhaustion signature of their transcriptome. By functional validation in a mouse model, we show how one such signaling axis, CCL20/CCR6, participates sufficiently in immune suppression that impairing it relieves the exhaustion phenotype and changes animal survival. This is particularly important because it raises the general prospect that inhibiting activating signals can resolve cancer immune suppression.

Identification of the CCL20/CCR6 axis as relevant for tumor control provides other important information with medical translation potential. This axis is implicated in a number of inflammatory and immune activated states including autoimmune disease and cutaneous T cell lymphoma. The potential for modulating the axis to reduce the activated states of immune cells and tumor cells has been extensively explored and led to early stage clinical trials (Getschman et al., 2017; Robert et al., 2017). Specifically, anti-CCL20 was advanced to clinical testing with a focus on inflammatory disease (Bouma et al., 2017; Laffan et al., 2020). However, the drug did not advance in that setting (Hippe et al., 2020). The data presented here suggest that it may have potential, not as a direct suppressor of cancer cell growth as proposed previously, but as a means of relieving immune exhaustion (Lee et al., 2017; Lu et al., 2017; Walch-Ruckheim et al., 2015). This would be of importance in the setting of bone metastases and prostate bone metastases in particular as these have shown little responsiveness to ‘check-point’ blockade (Goswami et al., 2016).

The tumor-specific myeloid populations (TIM and TAM) are the likely sources of the CCL20 ligand in the patient samples. CCL20 is expressed by both TIM and TAM populations. In TIM, the expression levels of CCL20 are strongly correlated with the overall TIM abundance (Fig. 7H, S7G). The distinctive TIM expression signature we identified in prostate bone metastases compared with other cancer types does raise the possibility that the CCL20/CCR6 axis is particularly important in prostate cancer. However, studies of other cancer metastasis, in the bone and other sites will be needed to deconvolve the contributions of tumor-intrinsic properties and the host tissue. It will also be important to consider systemic factors like androgen deprivation. Androgen deprivation therapy is universally applied to all prostate cancer patients as part of their standard treatment, and will require examination of rare outlier cases to define its role versus prostate cancer itself in changing CCL20/CCR6. We hope the data presented here will provide a foundational resource for further exploring this and other signaling axes and cellular relationships that inhibit immune response or remodel the bone marrow resulting in a tumor-permissive environment. Defining new approaches to the devastating clinical problem of prostate cancer bone metastases is critically needed.

## Data availability

The expression datasets generated in this study are available through Gene Expression Omnibus with the accession number GSE143791 (Reviewer token: mhwvcmwmhtijtaj). Interactive views of the single cell datasets, differential expression results, code notebooks, cell annotation and RData objects are available on the author’s website at http://pklab.med.harvard.edu/bonemet/.

## Acknowledgements

We are particularly indebted to our patients and their clinical care teams. We gratefully acknowledge support from Bill & Cheryl Swanson and from Gunther & Maggie Buerman. We acknowledge funding from NIH CA193481 and DK103074 (to DTS and PVK), National Cancer Institute CA 163191 (to DTS), HL131768 (to PVK), Dana-Farber / Harvard Cancer Center Nodal Award (CCSG grant P30CA006516), the Harvard Ludwig Cancer Center, the Harvard Stem Cell Institute and the Gerald and Darlene Jordan Professor of Medicine Chair (to DTS). N.B. was funded by the Swedish Cancer Society and Swedish Childhood Cancer Fund. N.B and K.G were funded by the Swedish Research Council. N.S was a recipient of the AACR-Millennium Fellowship in Prostate Cancer Research. Y.K was supported by a grant from the STARR cancer consortium. The Benign bone marrow material was provided by Mark A Randolph from the plastic surgery research lab at Massachusetts General Hospital. Olga Kharchenko designed the medical illustration in Figure 1A and the graphical abstract. Patient samples were sorted at the HSCI/CRM flow cytometry core facility at MGH. We thank the Center for Skeletal Research at Massachusetts General Hospital, supported by a grant from NIAMS (P30 AR075042) at NIH.

## Author contributions

N.B., Y.K., N.S., D.B.S. and P.J.S. conceived the study. P.J.S. coordinated the multi-disciplinary teams and the IRB-approved protocol. N.B., N.S., Y.K., P.S., J.H.S., D.B.S., D.T.S. and P.V.K. directed the study. N.B., Y.K., N.S., K.G. and T.H. designed experiments. Sample collection methodology and surgeries were performed by J.H.S. Human malignant samples were collected, isolated and libraries prepared by N.B. Y.K. collected human benign samples. FACS was performed by Y.K. Flow cytometry was performed by Y.K., T.H. and K.G., T.H. and K.G. collected human samples for flow cytometry analysis. ELISA measurements were performed by T.H. N.S. established the bone metastatic mouse model. Y.K., N.S., K.G., and T.H. performed in vivo experiments. T.H. and K.G. performed FACS and library preparations on samples from mouse experiments. S.M. and P.V.K. performed the computational analysis. N.B., Y.K., N.S., S.M., T.H., K.G., J.H.S., P.J.S., D.T.S., D.B.S. and P.V.K. interpreted the data and wrote the manuscript. All authors read, edited and approved the manuscript.

## Competing interests

P.V.K. serves on the Scientific Advisory Board to Celsius Therapeutics Inc. D.T.S is a director and shareholder for Agios Therapeutics and Editas Medicines; a founder, director, shareholder and scientific advisory board member for Magenta Therapeutics and LifeVault Bio, a shareholder and founder of Fate Therapeutics, and a director, founder, shareholder for Clear Creek Bio, a consultant for FOG Pharma and VCanBio and a recipient of sponsored research funding from Novartis. D.B.S. is a founder, consultant and shareholder for Clear Creek Bio.

## Methods

### Patient material

All human-subjects tissue collection was carried out with institutional review board (IRB) approval (Dana Farber/Harvard Cancer Center protocol 13-416 and Partners protocol 2017P000635/PHS).

### Surgical approach and collection of tumor and bone marrow specimens

#### Tumor specimens

In each case, the patient was brought to the operating room for clinically indicated decompression and stabilization in the setting of spinal cord compression related to metastatic prostate cancer. Each patient consented to use of their tissue for research purposes. Bone marrow (*e*.*g. Involved*) and tumor samples were taken with the patient in the prone position under general anesthesia as the spine was approached posteriorly. After the levels of the spine were exposed and identified, a Jamshidi needle was malleted directly into the desired vertebral body and a syringe connected to the needle was used to aspirate bone marrow immediately upon cannulation. Cannulation of the vertebral body is standard prior to placing stabilizing instrumentation into the bone. By placing the Jamshidi needle directly into the bone, this ensures that the aspirate from deep within the vertebral body is not diluted by surrounding blood in the surgical field or surgical irrigation. The marrow aspirate taken from the vertebral body is then directly stored into collection tubes and transferred to the laboratory for further sample preparation. Similar technique was utilized for the distant vertebral body level samples (*e*.*g. Distal*). During each surgery, several vertebral body levels above and below the primary site of spinal cord compression are instrumented for stabilization. As such, there is access to numerous vertebral body levels through a single surgical approach. After the vertebral body marrow had been collected and the spine instrumented, the spinal cord is decompressed through a laminectomy. Epidural tumor that is circumferentially surrounding and compressing the spinal cord is taken directly from the field and transferred to the laboratory for further processing. Specimens were also submitted to pathology for standard confirmation of diagnosis of metastatic prostate cancer.

### Animals

CCR6-KO mouse models were developed by Deltagen, Inc and ordered from The Jackson Laboratory and (B6.129P2-*Ccr6*^*tm1Dgen*^/J, #005793) and compared to age and gender match control mice C57BL6/J mice (#000664). All mice were maintained in pathogen-free conditions and all procedures were approved by the institutional Animal Care and Use Committee of Massachusetts General Hospital.

Mice with signs of bone metastasis, as shown by bioluminescence imaging, were included in the survival study during which the health condition of the mice was scored. We adapted this scoring method from Nunamaker et al., to refine the endpoint for mice bearing bone metastasis to minimize the pain and distress associated with bone metastasis progression (Nunamaker et al., 2013). For each category, mice are scored from 0 to 3: Body posture, eye appearance and activity level. Mice with a score at 3 in one of the categories or a cumulative score >6 are euthanized. The health was assessed by the animal facility veterinarian and the lead investigator for the specific experiment. The main cause of euthanasia were eye lesions due to metastatic burden in the mandible. In addition, some of the mice developed leg paralysis. Otherwise, a general poor body condition was observed in moribund mice. Only mice showing signs of metastatic tumors in the bones by bioluminescence imaging were included in the survival studies. For inclusion of mice for survival studies, we image mice with bioluminescence imaging at week 1, 2 and 3, to ensure that the mice that are included in the study were continuing showing exponential growth.

### Development of syngeneic prostate bone metastases cell line

The RM1-BoM1 cell line was obtained from the Power laboratory (Power et al., 2009) and was established from the parental prostate cancer mouse model RM1 (RAS and c-Myc oncogenes were overexpressed in normal epithelial prostate cell and injected in syngeneic mice to form tumors) (Power et al., 2009). We enriched RM1-BoM1 cells negative for GFP and transduced with Tdtomato (LeGO-T2, Addgene #27342) and luciferase (pENTR-LUC, Addgene, #17473). We injected 2.10^5^ RM1-BoM1 cells into the left ventricle of C57Bl/6 mice and monitored bone metastases development by sequential bioluminescence imaging. Bone metastatic cells were harvested and selected for the Luciferase and Tdtomato expression. We used the mouse as a bioreactor and repeated a second round of injection and isolation of bone metastases cells in order to establish the RM1-BoM3 bone metastatic cells. The RM1-BoM3 sells were maintained in DMEM (Corning, 15-013-CV) complemented with 10% FBS (GIBCO by Life Technologies, A31605-01) and 1% Penicillin-Streptomycin (GIBCO by Life Technologies, 15140-122).

### In-vivo Bioluminescence imaging

Mice were anesthetized by isoflurane inhalation with 2% O_2_ and 4.5mg/mouse of D-luciferin K salt (RR labs Inc., San Diego) was administered by intraperitoneal injection. After 5 minutes, mice are imaged using a SPECTRAL Ami X allowing to detect the localization of the cancer cells in the mice and to measure the luciferase activity.

### CCL20 blocking antibody treatment

C57BL6/J male mice (#000664) received an intraperitoneal injection of 45μg anti-CCL20 blocking antibody (R&D Systems clone 114908) or rat IgG isotype control antibody (R&D Systems, clone 43414) 1 day prior to intracardiac injection of RM1-BoM3 prostate cancer bone metastasis cells. The administration of blocking antibody and isotype control then continued on a twice-weekly basis until the end of the experiment.

### Sample preparation

#### Dissociation of tissues into single cells

All samples were collected in Media 199 supplemented with 2% (v/v) FBS. Single cell suspensions of the tumors were obtained by cutting the tumor in to small pieces (1mm^3^) followed by enzymatic dissociation for 45 minutes at 37°C with shaking at 120 rpm using Collagenase I, Collagenase II, Collagenase III, Collagenase IV (all at a concentration of 1 mg/ml) and Dispase (2mg/ml) in the presence of RNase inhibitors (RNasin (Promega) and RNase OUT (Invitrogen). Erythrocytes were subsequently removed by ACK Lysing buffer (Quality Biological) and cells resuspended in Media 199 supplemented with 2% (v/v) FBS for further analysis.

#### ELISA measurement

CCL20 protein levels measured in plasma collected from bone marrow of bone-metastatic prostate cancer patients (*Involved, Distal*) and from bone marrow of patients undergoing hip replacement surgeries (benign BM) using a commercially available enzyme-linked immunosorbent assay (ELISA kit R&D Systems, Minneapolis, USA) according to the manufacturer’s protocol. Absorbance was measured with Synergy HTX multi-mode reader (Bio-Tek).

#### Bone marrow processing

Bone marrow samples were filtered using 70 micron filter then centrifuged at 600 g for 7 minutes at 4°C. Plasma were collected followed by erythrocytes removal using ACK Lysing buffer (Quality Biological). Cells were resuspended in Media 199 supplemented with 2% (v/v) FBS for further analysis.

#### FACS sorting of human samples for single-cell RNA-sequencing

Single cells from tumor and bone marrow samples subjected to RBC lysis were surface stained with anti-CD235-PE (Biolegend) for 30 min at 4°C. Cells were washed twice with 2% FBS-PBS (v/v) followed by DAPI staining (1 ug/ml). Flow sorting for live and non-erythroid cells (DAPI-neg/CD235-neg) was performed on a BD FACS Aria III equipped with a 100um nozzle (BD Biosciences, San Jose, CA) instrument. All flow cytometry data were analyzed using FlowJo software (Treestar, San Carlos, CA).

#### FACS sorting of murine samples

In order to obtain bone marrow T cells and myeloid cells for single-cell RNA-sequencing from tumor bearing mice, RBCs were lysed and samples were subsequently incubated with anti-mouse Fc block (BD Pharmingen, 553142) for 10 minutes at 4°C. This was followed by surface staining for CD11b (myeloid cells) and CD3e (T lymphocytes) for 30 minutes at 4°C. The samples were then washed with 2% FBS-PBS (v/v) followed by resuspension in 2% FBS-PBS with 0.1μg 7-AAD. Flow sorting for live myeloid and T cells (7AAD-CD11b+ or CD3e+) was performed on a BD FACS Aria III equipped with a 70*μ*m nozzle.

For evaluation of CTL proliferation response in mice with bone metastasis, bone marrow was stained with 3*μ*M CellTrace CFSE (Thermo Fisher Scientific) in accordance with the manufacturer’s instructions. This was followed by a 10 min incubation with anti-mouse Fc block and a 30 minute cell surface staining with anti-mouse CD4-APC/Cy7 and CD8-PE/Cy7 (Both from Biolegend). CFSE labeled CD8+ CTLs were subsequently sorted on a BD FACS Aria III (BD Biosciences, San Jose, CA) equipped with a 70*μ*m nozzle.

#### FACS analysis

Independent samples from patients with prostate bone metastases were used for FACS analysis. Cells from human bone marrow and tumor samples were surface stained with lymphoid, myeloid and hematopoietic stem and progenitor antibody panels after blocking with anti human fc block (BD Pharmingen 564219) for 10 minutes at room temperature (Supplementary Table 7) for 30 min at 4°C. Cells were washed twice with 2% FBS-PBS (v/v). Samples that were to eventually be fixed and permeabilized were stained with LIVE/DEAD fixable viability dye (Thermo Fisher Scientific, Waltham, MA). For the intracellular staining, to determine Treg infiltration, cells stained with lymphoid surface markers were fixed and permeabilized with Cytofix/Cytoperm (BD Biosciences, San Jose, CA) for 20 min at 4°C. Cells were then washed twice with 1× Perm/Wash buffer (BD Biosciences, San Jose, CA) and incubated overnight at 4°C with anti-FoxP3-PE. On the following day, cells were washed twice in Perm/Wash buffer and resuspended in 2% FBS-PBS (v/v) for analysis.

For analysis of CTL exhaustion in bone metastasis bearing mice, bone marrow cells were blocked with anti-mouse Fc block (BD Pharmingen, 553142) followed by surface staining with the murine T cell exhaustion panel (Supplementary Table 7) for 30 min at 4°C. Cells were washed with 2% FBS-PBS (v/v) followed by resuspension in 2% FBS-PBS with 0.1μg 7-AAD.

Flow cytometry was performed on a BD FACS Aria III (BD Biosciences, San Jose, CA) instrument. All flow cytometry data were analyzed using FlowJo software (Treestar, San Carlos, CA) and Prism software. Statistical analyses were performed as indicated and *P* values of ≤0.05 considered significant.

#### Healthy bone marrow data

To provide an additional comparison to the samples from cancer patients and patients with benign inflammation that were collected as part of this study, we have also analyzed single-cell RNA-seq data for healthy individuals published by Oetjen *et al*. (Oetjen et al., 2018) The data was downloaded from GEO (GSE120221, GSE120446).

#### Massively parallel single cell RNA-sequencing

Single cells were encapsulated into emulsion droplets using Chromium Controller (10x Genomics). scRNA-seq libraries were constructed using Chromium Single Cell 3’ v2 Reagent Kit according to the manufacturer’s protocol. Briefly, post sorting sample volume was decreased and cells were examined under a microscope and counted with a hemocytometer. Cells were then loaded in each channel with a target output of ∼4,000 cells. Reverse transcription and library preparation was done on C1000 Touch Thermal cycler with 96-Deep Well Reaction Module (Bio-Rad). Amplified cDNA and final libraries were evaluated on an Agilent BioAnalyzer using a High Sensitivity DNA Kit (Agilent Technologies). Individual libraries were diluted to 4nM and pooled for sequencing. Pools were sequenced with 75 cycle run kits (26bp Read1, 8bp Index1 and 55bp Read2) on the NextSeq 500 Sequencing System (Illumina) to ∼70-80% saturation level.

### Analysis

#### Quality control and preprocessing of single-cell RNA-seq data

FASTQ files were processed with the CellRanger software (10x Genomics, Inc., version 2.0). Human genome hg19 was used as the reference genome (10x Genomics, Inc.) to generate the matrix files containing cell barcodes and transcript counts. Only cells with total UMI exceeding 600 were included in the downstream analysis. Statistics on the sequencing results are available in Supplementary Table 2. Samples were initially analyzed using Pagoda2 (https://github.com/hms-dbmi/pagoda2) for quality control and data exploration.

#### Joint clustering and cell annotation

We used Conos (https://github.com/hms-dbmi/conos) to integrate multiple scRNA-seq datasets together and to align our data with other public scRNA-seq data, closely following the Conos tutorial. Sample pre-processing was performed using default pagoda2 settings (“basicP2proc()” function). The integration was performed using Conos with *k* = 30, *k*_*self*_ = 15, CPCA space, nPC=50, and n.odgenes=500, with an angular distance measure. Louvain clustering was used to build to determine joint cell clusters across the entire dataset collection. The annotation of clusters was performed manually based on the marker genes, with some clusters being assigned same label. One positioned between T and B lymphocytes was determined to represent doublets and was omitted from the analysis. As an additional check the annotation was compared with published scRNA-seq data from healthy bone marrow (Oetjen et al., 2018). To do so, the raw count matrix was downloaded, and integrated into the joint graph using the same Conos settings. Conos label transfer was used to examine the correspondence of the annotation mapping. A 2D embedding (Fig. 1E) was generated using largeVis using default settings. To create a more detailed annotation of the T lymphocytes, we extracted all myeloid and all T cell populations (CD8+ T cells, CD4+ T cells, NK and NKT cells), and realigned separately using Conos. Leiden community detection method (as implemented in Conos) was used to determine refined joint clusters, providing higher resolution than the initial analysis. To generate an embedding for the T cell populations (Fig. 6A), a largeVis embedding was first generated in 13 dimensions using default parameters, and then reduced to a 2D embedding using t-SNE (using default perplexity value of 50). Analogous procedure was performed for the myeloid cells, however the myeloid subpopulations were visualized using the original 2D embedding that was generated for the entire dataset (Fig. 1E).

#### Analysis of cell proportions and compositional data analysis

Statistical significance of proportion differences (Fig. 3B, S3A) was evaluated using Wilcoxon rank sum test, using 1000 bootstrap resampling rounds to report p-values at the 99% reproducibility power (*i*.*e*. reporting 0.99^th^ quantile of the sampled p-values). The p-values in the figures were reported using the following symbols: *p<0.05, **p<0.01, ***p<0.001, ****p<0.0001.

As an alternative to the simple proportion tests, we also used compositional data analysis (Pawlowsky-Glahn and Buccianti, 2011) to analyze the differences between Tumor/Involved and Benign samples (Fig. S3D). The CoDA avoids the non-independence of cell type fractions (in each sample they sum up to one). Isometric log-ratio (ilr) transformation was first applied, which allows to work with (n-1) independent artificial variables (ilr coordinates/variables) instead of cell type fractions constrained in the n-dimensional Simplex space. Each ilr variable is a linear combination of logarithms of cell type fractions. The set of (n-1) ilr coordinates forms the Euclidean space, where standard methods to discriminate groups of samples are permitted. To separate groups of samples (*e*.*g*., Benign and Tumor) in the ilr-space, the canonical discriminant analysis was applied (using candisc package). The first canonical coordinate is a linear combination of log-fractions and represents a weighted contrast (balance) between cell types of positive and negative contributions to the linear combination. The loadings of different cell types on the first canonical coordinate were taken to be the separating coefficients (x axis, Fig. S3D). To evaluate the robustness and statistical significance of the separating coefficients, we performed random cell subsampling, repeating the procedure above 1000 sampling rounds. In each subsampling round, 500 cells were randomly sampled from each dataset to account for limited detection rates of cell types across datasets. Furthermore, to account for inter-patient variability, in each sampling round we performed bootstrap sampling of the datasets. Then the ilr calculation and subsequent procedures were repeated, and the separating coefficients resulting from different sampling rounds were shown on Fig. S3D as boxplots.

#### Analysis of expression distances

Expression differences between matching subpopulations (Fig. 3C, S3B) were determined by first estimating “mini-bulk” (or meta-cell) RNA-seq measurements for each subpopulation in each sample. Briefly, in each dataset, the molecules from all cells belonging to a given subpopulation were summed for each gene (*i*.*e*. forgetting cellular barcodes). The distance between the resulting high-coverage RNA-seq vectors was calculated using Jensen-Shannon divergence (JS). Expression distances between samples (Fig. 3C, S3B) were determined as a normalized weighted sum of JS divergence distances across all cell subpopulations contained in both samples, with the weight equal to the subpopulation proportion (measured as a minimal proportion that the given cell subpopulation represents among the two samples being compared).

Cell-specific magnitude of expression shifts between fractions ***XX*** and ***YY*** for the individual cell types (Fig. S3C) was determined by normalizing between-fraction distances by the mean within-fraction distance. Specifically, for samples *i, j* such that *i ∈* ***XX*** and *j ∈* ***YY***, standardized distance 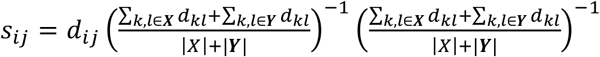, where *d*_*ij*_ is the angular distance of the log-normalized mini-bulk molecule counts for the samples *i* and *j*. The boxplots in Fig. S3C show the *s*_*ij*_ values for the different classes of sample pairs. A minimal number of 10 cells (of the selected cell type) were required for a sample to be included in the comparison.

#### InferCNV analysis (Figure S2A)

To characterize copy number aberrations from the single cell RNA-seq data, inferCNV (Patel et al., 2014) (https://github.com/broadinstitute/inferCNV) was used. The raw count matrices for the measured tumor cells were extracted from the Conos object. For control cells, we have used normal epithelial prostate cells from three healthy prostate samples, published by Henry *et al*. (Henry et al., 2018) The count matrices for the control data were downloaded from GEO under the accession number GSE120716. For each of the three patients we randomly selected 300 epithelial cells to serve as a control. The following inferCNV parameters were used: denoise=TRUE, cutoff=0.1.

#### Analysis of intra-tumoral heterogeneity (Figures 2, S2)

Only samples with more than 10 annotated tumor cells were considered in the analysis. The datasets were normalized using pagoda2 (trim=0, min.n.odgenes=1000, nPcs=3). To examine to what extent the first principal component (PC) of each datasets separates cells in other datasets, the cells of each sample were projected onto the first PC of each of the samples (Fig. S2F). Size-normalized log counts were used for the projection, without taking into account the dataset-specific variance normalization. Tumor cell datasets were integrated using Conos (k=10, k.self=5, ncomps=10, metric=’L2’, n.odgenes=1000), and a joint largeVis embedding was obtained for visualization. Cell clusters were determined using Leiden community detection algorithm with resolution of 0.6.

#### Expression signature scores

To account for the complex gene expression patterns in M2 tumor associated macrophage, monocytes differentiation to macrophage, Treg activity, CD8+ T cell dysfunction and cytotoxicity, The signature scores were estimated as average expression values of the genes in a given set. Specifically, we first calculated signature score for each cell as an average normalized (for cell size) gene expression magnitudes, then the signature score for each sample was computed as the mean across all cells. All signature gene modules are listed in the Supplementary Table 4. The M2 signature genes were from Aziz *et al*. (Azizi et al., 2018). CD8+ T cell cytotoxicity were measured by expression of CD8A, CD8B, GZMA, GZMM, GZMB, GZMK, GZMB, PRF1 (Rooney et al., 2015). The CD8+ T cell dysfunction genes were taken from Li *et al*. (Li et al., 2019). To define gene signature of monocytes to macrophage differentiation, we took top 100 differentially expressed genes based on published *in vitro* studies of monocytes to macrophage differentiation (Liu et al., 2008; Martinez et al., 2006). For this, the microarray data was analyzed by using affy (Liu et al., 2008) R package, and the differentially expressed genes were identified using limma (Martinez et al., 2006) R package. The statistical significance was assessed using Wilcoxon rank-sum test. To ensure robustness of the result, two types of bootstrap resampling were performed: i) resampling cells, and ii) resampling genes. 1000 bootstrap resampling rounds were formed for each scenario, and a p-value corresponding to the 99% reproducibility power was reported in the figures using the following symbols: *p<0.05, **p<0.01, ***p<0.001, ****p<0.0001.

#### Differentially expressed genes

Differential expression (DE) and marker gene detection was performed using Wilcoxon rank sum test, implemented by the getDifferentialGenes() function from Conos R package. The genes were considered differentially expressed if the p-value determined Z score was greater than 3. Since the genes were used primarily for pathway enrichment analysis, the Z score was not corrected for multiple hypothesis testing. The getDifferentialGenes() function was used to identify differences between similar subpopulations (CTL-1 vs. CTL-2; Mono1 vs. Mono2 vs. Mono3).

For differential expression analysis between sample fractions (for example *Tumor* Treg vs. *Benign* Treg), getPerCellTypeDE() function in Conos was utilized. As described previously, it first forms “mini-bulk” (or meta-cell) RNA-seq measurements by combining all molecules measured for each gene in each subpopulation in each sample. This results in a collection of bulk-like RNA-seq samples, and the differential expression problem is then reformulated as a standard bulk RNA-seq differential expression problem DESeq2 to compare these “bulk-like” meta-cell samples, using appropriate design models (i.e. *Tumor* vs. *Benign* factor on samples). This approach is suitable for the comping groups of patient samples, as it focuses on the inter-individual variability (variability between patients) as opposed to cell-to-cell variability within each sample which is much smaller. At the same time, the overall depth of the “mini-bulk” profiles enable to estimate the uncertainty of each the expression magnitudes (*i*.*e*. the samples with large number of cells will result in mini-bulk profiles of greater depth). Preliminary benchmarking shows that such ‘mini-bulk’ approach is effective for differential expression testing (Crowell, 2020). A minimal number of 10 cells (of the selected cell type) were required for a sample to be included in the comparison. The DESeq2 analyses were ran using “∼fraction” model. For paired tests (within individual), “∼patient+fraction” model was used. The p-values were translated into Z scores, with positive scores corresponding to upregulation, negative to downregulation. Several additional tests were implemented in additional to the mini-bulk based tests described above: i) a Wilcoxon rank sum test across mini-bulk samples (wZ); ii) Wilcoxon rank sum test on samples with cells resampled using bootstrap sampling (100 sampling rounds), reporting the Z score at 90% reproducibility power (bwZ); and iii) Wilcoxon rank sum test across single-cell expression matrix (i.e. without collapsing cells into “mini-bulk” samples) (cwZ). These tests are included in the interactive differential expression tables on the author’s website.

#### Comparative analysis with public cancer datasets (Figure 5)

To compare the profile of myeloid and T lymphocyte populations observed in our bone marrow study with the immune microenvironment of other cancer types, we used *Conos* (with *k* = 30, *k*_*self*_ = 10 CPCA rotation space and an angular distance measure) to perform a joint alignment of myeloid cells from benign bone marrow (Oetjen et al., 2018), prostate cancer bone metastasis, pancreatic cancer, lung cancer (Lambrechts et al., 2018) and liver cancer (Zhang et al., 2019) samples. Bone marrow cell annotations were propagated to other cancer types using *propagateLabels()* function in *Conos*. For myeloid cells, we measured the cell proportion of TIM and TAM across cancer types, compared monocyte inflammatory signature score for the TIM-like cells, and the M2 macrophage score for the TAM-like. Select marker genes were shown in the heatmaps (Fig. 5E,F). For comparison of the T lymphocyte states, only study containing matched (adjacent) normal control samples were analyzed. Joint alignment and label propagation were then performed in the same way as for monocytes. We then evaluated T cells exhaustion scores in CTLs, observing the expected increase in exhaustion signature between the tumor and adjacent normal samples, despite variable base-level expression level of between the tissues (Fig. S6I).

#### Ligand and Receptor analysis (Figure 6H,I)

To screen for potential channels of communications between different cell types, we looked at expression of previously annotated reciprocal ligand-receptor pairs. The annotated list containing 1307 pairs of well-annotated receptors and ligands was downloaded from CellPhoneDB (Vento-Tormo et al., 2018).. Both ligands and corresponding receptors were required to be detected in more than 10% of the cells of a given type. Initial filtering reveled 241 ligand-receptor pairs between myeloid cell and T cells. (Fig. 6H, Supplementary Table 5). Further filtering for ligand-receptor channels potentially connecting TIM/TAM with T cell populations was performed by requiring ligands to be upregulated in TIM or TAM cells (compared to other myeloid cells, Zscore >5), and requiring the corresponding receptor to be upregulated (log fold change >0) in T cells in *Tumor* compared to *Benign* (Fig. S6I, Supplementary Table 5). Differentially expressed genes were identified using Wilcoxon rank sum test, implemented by the getDifferentialGenes function in Conos.

#### Survival analysis on bulk data (Figure S4E)

To test if a given gene signature is associated with differential survival of cancer patients (Cancer Genome Atlas Research et al., 2013) (Fig. S4E), we first calculated average expression of the signature in each cancer type based on the bulk RNA-seq data. The bulk patient samples were then stratified into two groups, based on average signature expression, separating patients with top 25% of scores and bottom 25%. A standard Kaplan-Meier survival analysis was then used to determine the association of these groups with survival rate. Kaplan-Meier survival analysis in Fig. 7C,D and Fig. S4E were performed using the *survival* R package.

#### Gene ontology and gene set enrichment analysis (Figures 3D,E S3E,F)

*clusterProfiler* package was used to evaluate enrichment of the GO BP categories in the sets of top 300 up- and down-regulated genes separately. The set of all expressed genes was used as a background. The categories with adjusted p-value of enrichment below 0.05 were then clustered into 20 clusters based on the similarity of the participating genes (*i*.*e*. the genes out of the 300 genes being tested that fall within each category, using binary distance measure). The clusters were named according to the most significantly enriched category, shortening some names to fit the plot.

**Figure S1.**
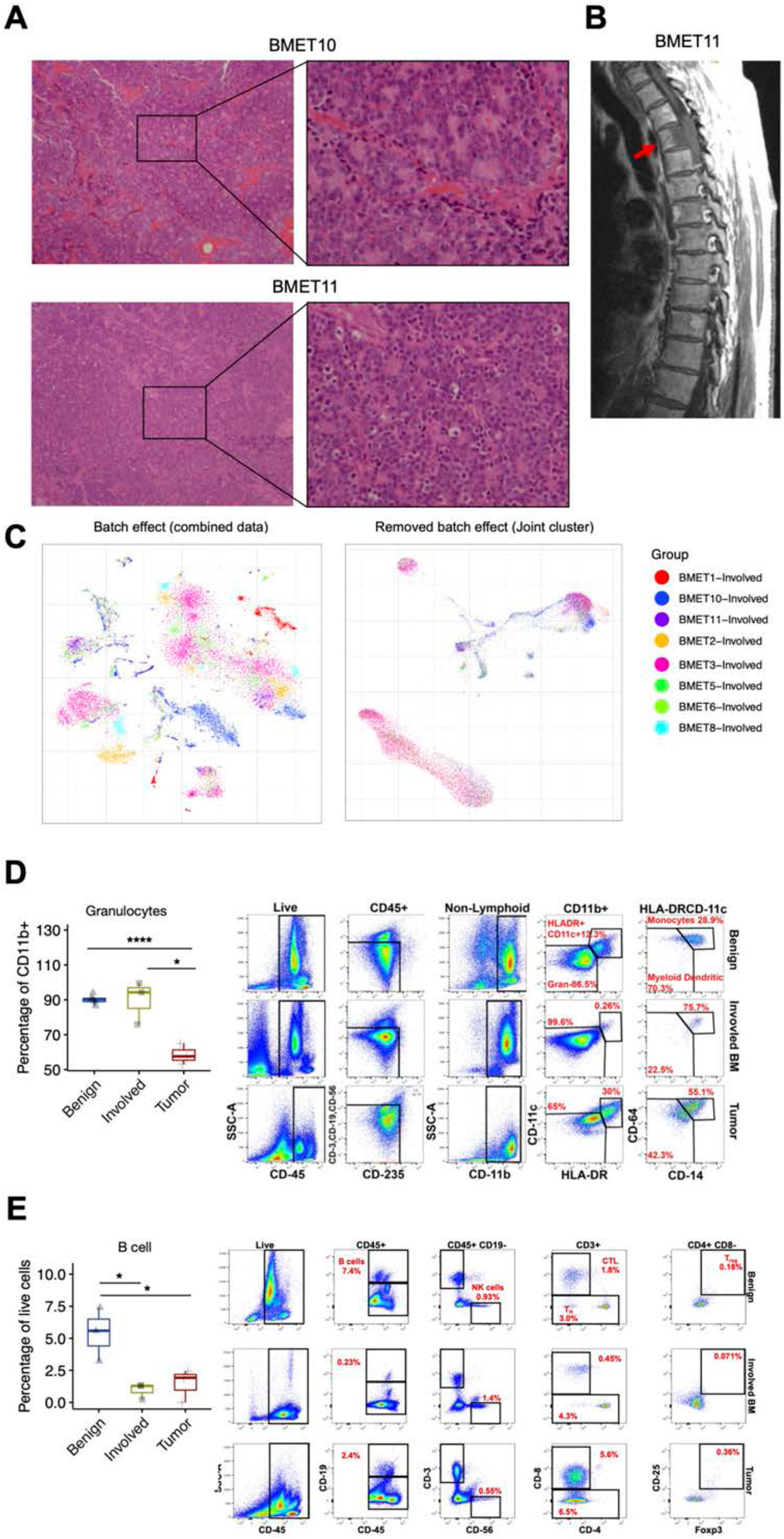
Sample properties and gating strategies. **A** H&E-stained tissue sections of metastatic tumor resected from two additional patients representative patients with bone metastatic prostate cancer (BMET10, BMET11) are shown as in Figure 1B. **B** Similar to Figure 1C, the sagittal T1 MRI imaging of the thoracic spine is shown for patient BMET11. An infiltrating soft tissue mass within the T3 vertebral body (arrow) with extension into the epidural space from T2 through T4. **C** An embedding combining the Involved BMET samples is shown without any corrections (left) and with Conos integration (right), illustrating strong inter-individual sample differences. **D** Boxplots showing flow cytometry analysis of granulocytes (Supplementary Table 7) proportion in Benign, Involvedand Tumor fractions (left) from three independent patients. Statistical analysis was performed using a Student’s t-test (*p<0.05; **** p < 0.0001). Gating strategy for flow validation of myeloid populations. Labels above the flow plots refer to the parent population and the percentages are of parent gate (right). **E** Boxplots showing flow cytometry analysis of **B** cells (Supplementary Table 7) proportion in Benign, Involved and Tumor fractions from three independent patients. Statistical analysis was performed using a Student’s t-test (*p<0.05). Gating strategy for flow validation of lymphoid cell types. Labels above the flow plots refer to the parent population in the percentages are of the parent gate (right).

**Figure S2.**
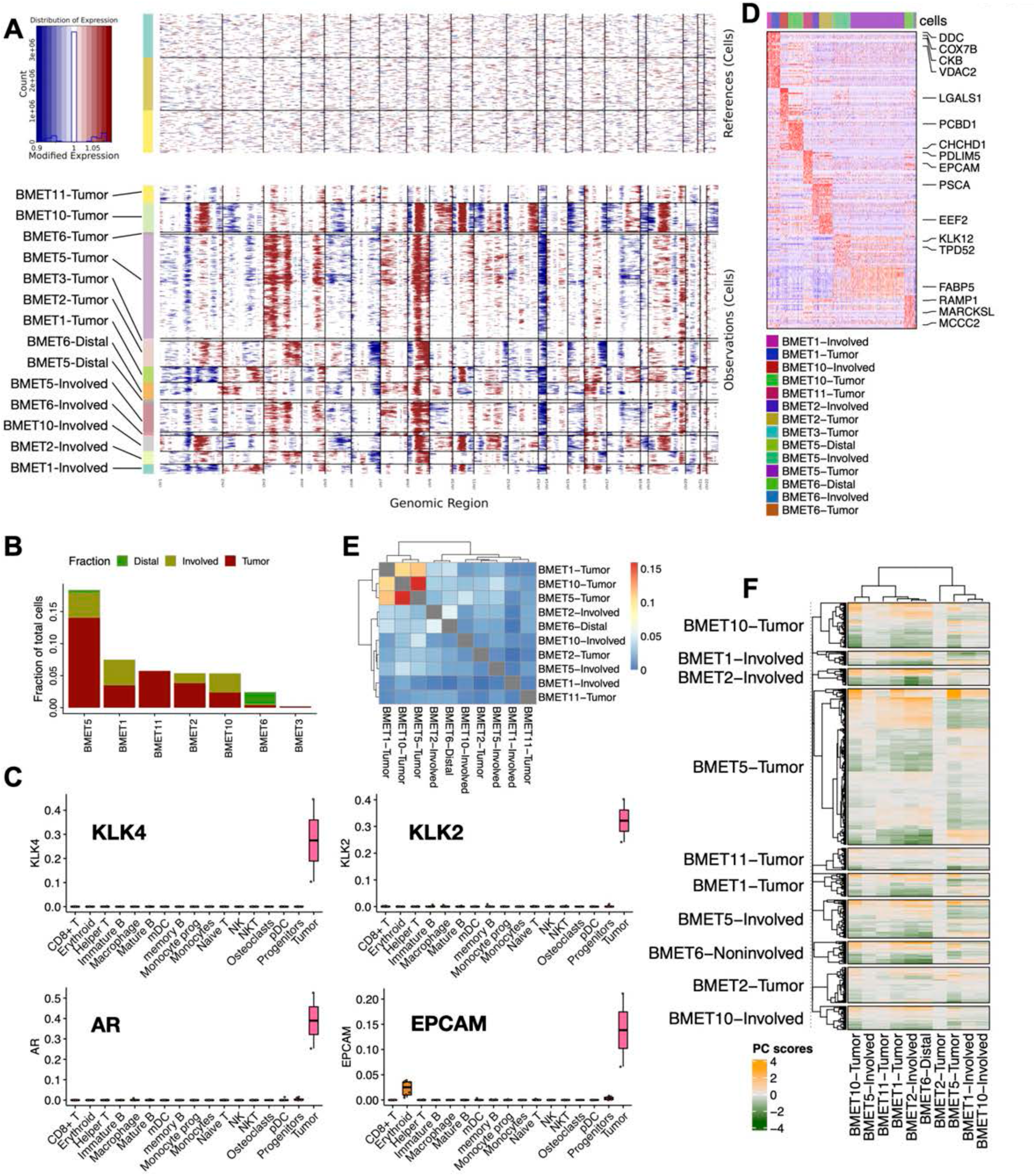
Characterization of tumor cells. **A**. lnferCNV analysis shows pronounced patient-specific CNV patterns in the tumor cells (bottom), relative to the healthy prostate epithelial cellcontrol (top) (see Methods). **B**. Number of tumor cells detected in Distal, Involved and Tumor fractions is shown for different patients. **C**. Boxplots show expression of the prostate cancer markers in the tumor cells detected in the *Distal* fractions. **D**. Heatmap shows an overview of genes (rows) differentially expressed between tumor cells of different patients. **E**. Overlap of top-scoring 200 genes in the PCs determined for the tumor populations in different samples is shown as a heatmap. The color shows proportion of overlapped gene. **F**. A heatmap showing cell-specific scores (rows) of cells in different samples (row blocks) for the first principal components (PCs) of intra-tumoral variation within different tumors (columns). The first PCs determined in one sample separate cells in other samples, suggesting common axes of variation.

**Figure S3.**
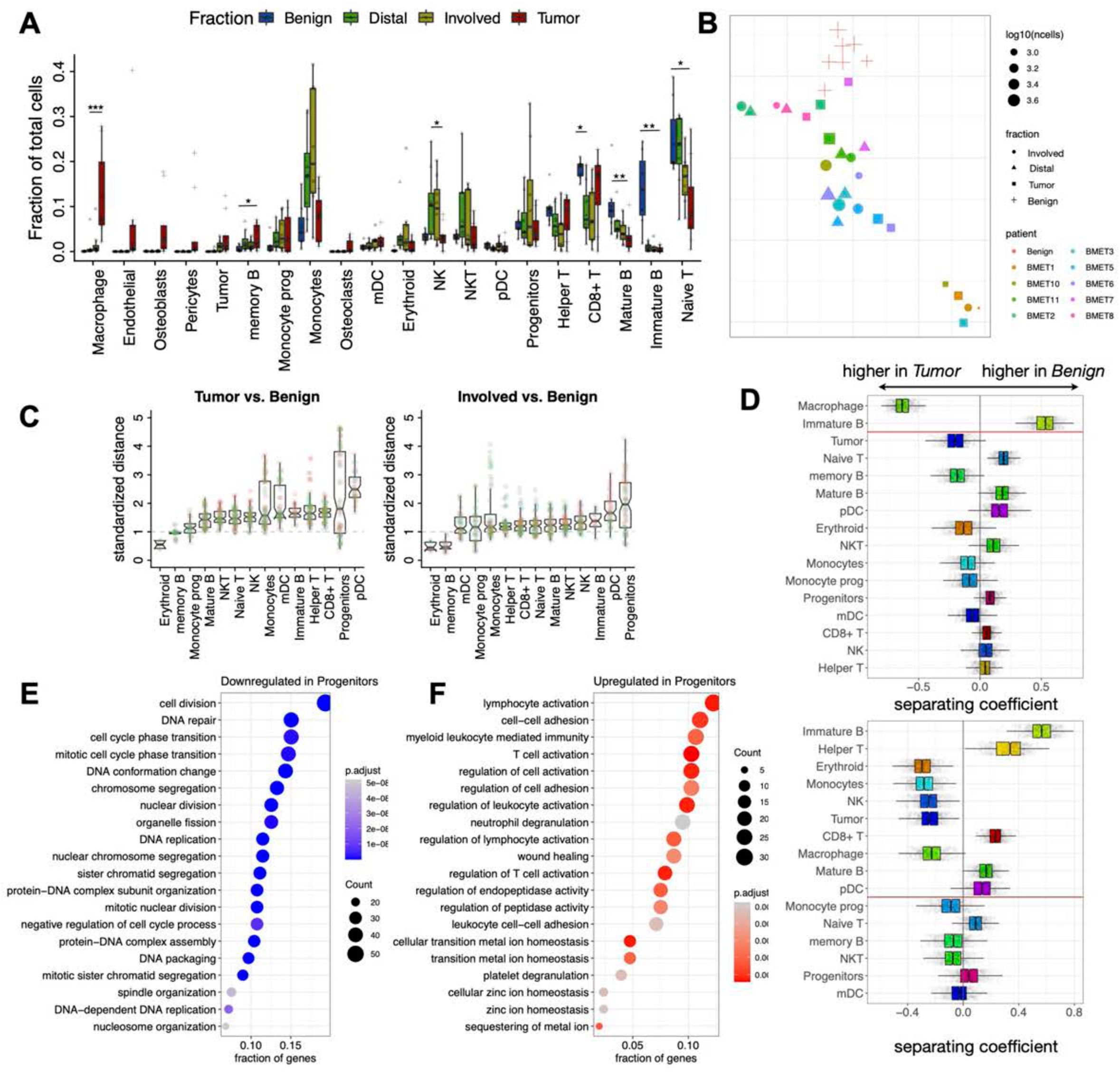
Overview of compositional differences between fractions. **A**. Similar to Fig. 3A, the boxplots compare relative abundance of different cell populations between Benign controls and the three cancer fractions for all major cell populations. **B**. At SNE embedding of different samples, based on their overall expression distance. The similarity measure measures the magnitude of expression change for each subpopulation, using size-weighted average to combine them into an overall expression distance that controls for compositional differences. Colors demarcate patients (all Benign controls are shown in one color), symbols indicate different sample fractions. **C**. The extent of expression differences is shown for four different contrasts. The boxplots show the expression distance of pairs from two distinct classes (i.e. Tumor vs. Benign), normalized by the median expression distance of all sample pairs within each class (see Methods). **D**. Contrast of cell type abundance proportions discriminating Benign and Tumor samples. The coefficients (x-axis) are loadings associated with each cell type separating the two types of samples in the isometric log-ratio simplex space. The boxplots show results of 1000 bootstrap cell resampling rounds (see Methods). f. Analogous contrast loadings for surfaces discriminatingbetween Benign and Involved samples. **E**,**F**. GO BM categories with most significant enrichment for the top 300 down-(E) and up-regulated (F) genes within the Progenitors population are shown for the Involved vs. Benign contrast.

**Figure S4.**
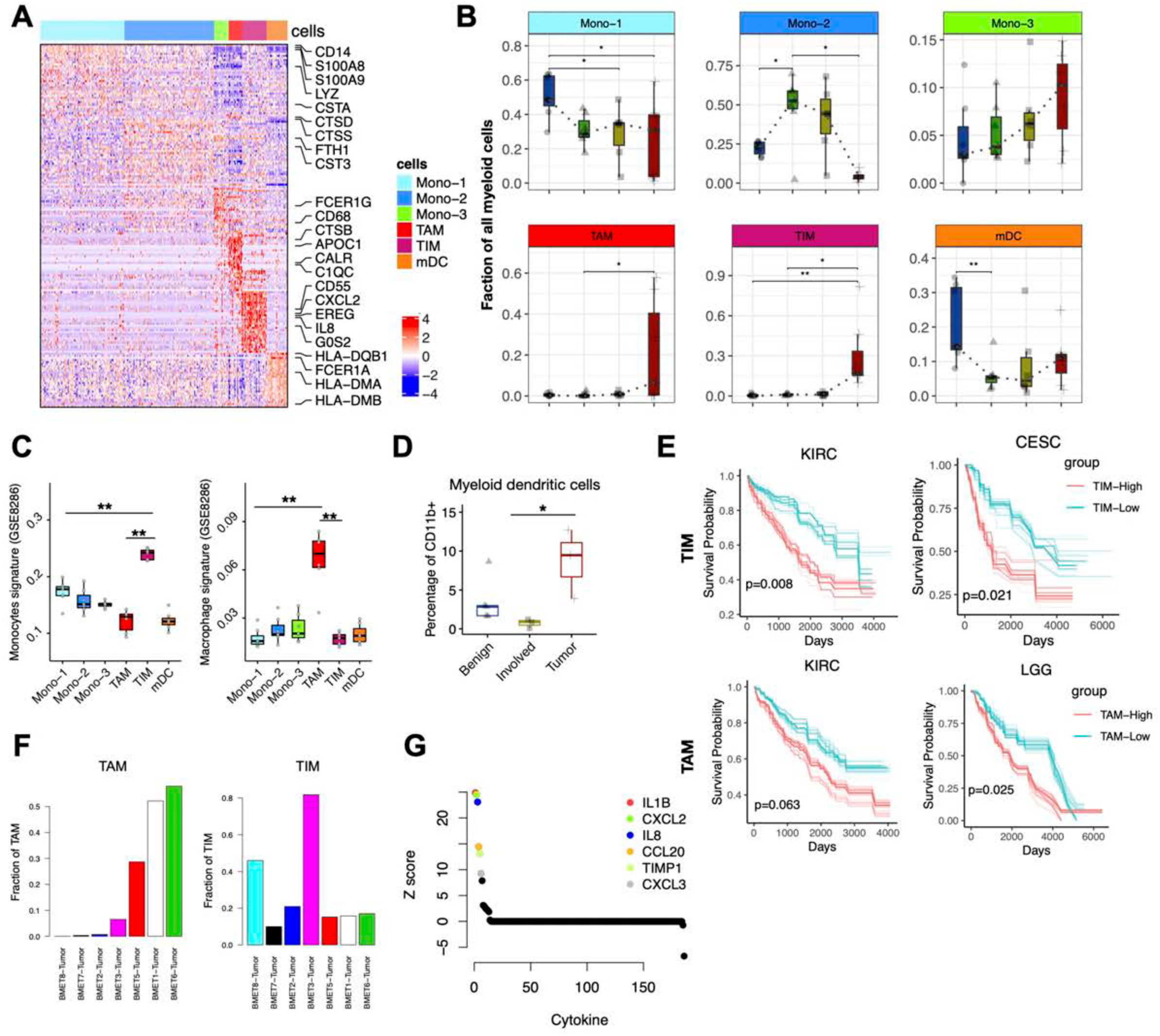
Myeloid populations shift towards inflammatory monocytes and repressive macrophages in the Tumor fraction. **A**. Heatmap of the highly expressed genes in different myeloid populations. **B**. Comparison of relative abundance of different myeloid cell clusters in *Benign, Distal, Involved* and *Tumor* fractions. **C**. Average expression of monocytes and macrophage signature genes selected based on in vitro differentiation of monocytes to macrophages (see Methods). **D**. Boxplot showing flow cytometry analysis of Myeloid dendritic cells (SupplementaryTable 7) in *Benign, Involved* and *Tumor* fractions from three independent patients. Statistical analysis was performed using a Studen’ts t-test (*p<0.05).**E**. Kaplan-Meier survival curves for TCGA cancer types: cervical squamous carcinoma (CESC), kidney renal clear cell carcinoma (KIRC), Low Grade Glioma (LGG). Patients were stratified into two groups based on the average expression (binary: top 25% versus bottom 25%) of TIM and TAM expression signatures as annotated by key marker genes in Supplementary Table 4. Bootstrap resampling were performed on signature genes and p-value was calculated using the 95% reproducibilitypower p-value. **F**. Relative abundance of TIM and TAM cells in different tumor samples, measured as a fraction of myeloid population in each sample. **G**. Cytokines ranked by the significance of differential expression (y axis) comparing TIM with all other cell types. The names of the 6 highest expressed TIM-specific cytokine genes are shown.

**Figure S6.**
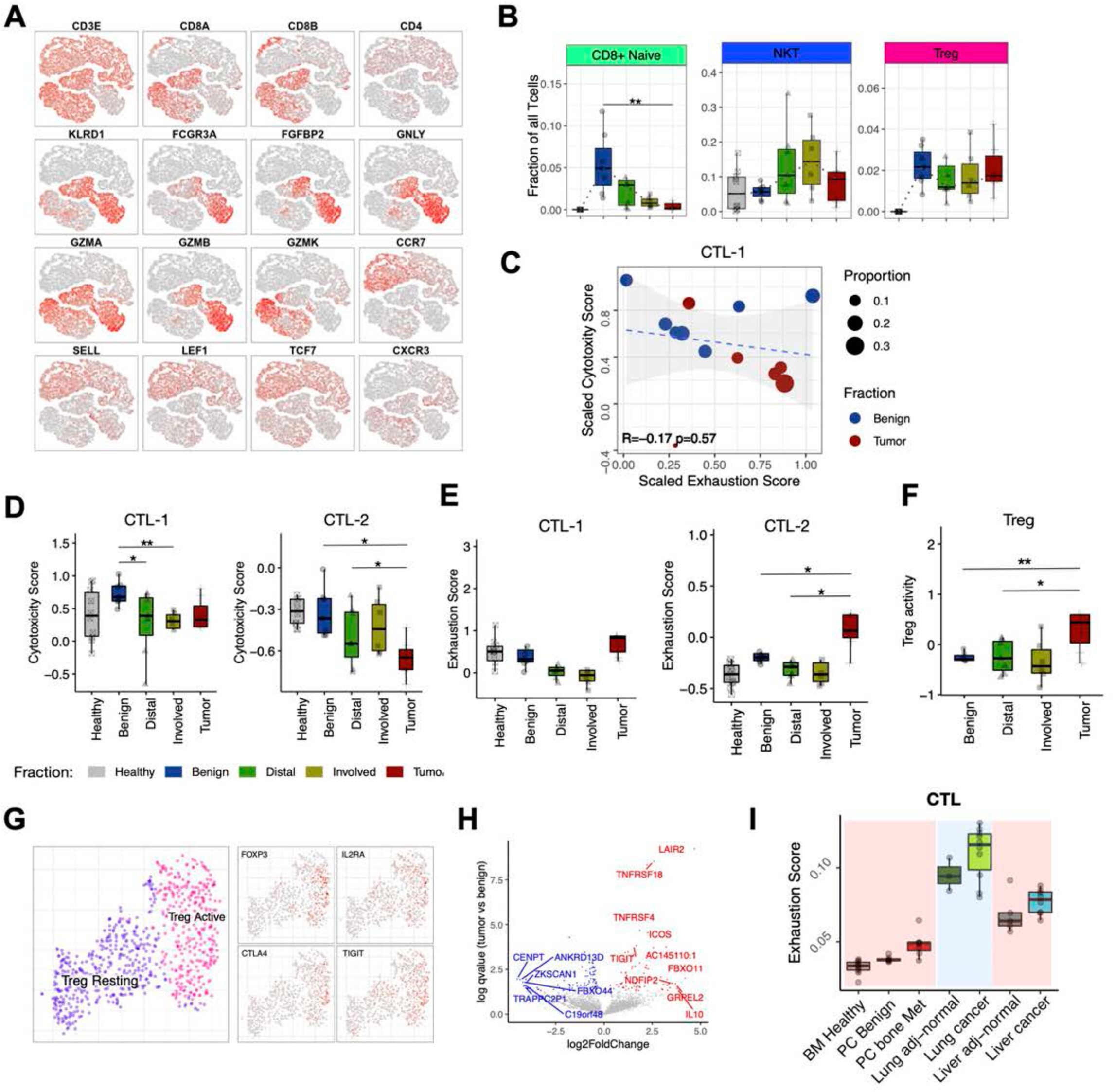
Tumor fraction shows increased abundance of exhausted cytotoxic populations and helper T cells. **A**. Expression of marker genes for different T cell subpopulations, shown on a joint T cell embedding. **B**. Comparison of proportions of CD8+ na”fve, NKT and Treg in in *Benign, Distal, Involved* and *Tumor* fractions. C. Similar to Figure 6E, the relationship between exhaustion and cytotoxicity is shown for the CTL-1 population. **D**. Cytotoxicity of CD8+ **T** cells. Boxplots illustrate significant decrease in the cytotoxicity of the CTL-2 subpopulation in the *Tumor* fraction. **E**. Exhaustion of CD8+ cytotoxic **T** cells. Boxplots show composite exhaustion scores of CTLs in different fractions, highlighting significant increase of exhaustion signature in the *Tumor* fraction. See Supplementary Table 4 for the genes defining exhaustion signature. **F**. Boxplots illustrate significant increase of Treg activity of Treg subpopulation in the *Tumor* fraction. Treg activity was measured as an average expression of genes shown in Supplementary Table **4. G**. Focused re-analysis of Treg compartment reveals activated and resting Treg subpopulations. Expression of key marker genes a visualized on the same tSNE embedding. **H**. Volcano plot showing differentially expressed genes between active Treg cells in *Benign* and *Tumor* fractions. The plot shows negative 10910 transformed Q values (y axis), as a function of the average109 2 fold changes in expression (x axis). The genes exhibiting significantly higher expression in *Tumor* faction are shown on the top (red), while those significantly higher in *Benign* fraction are shown on the bottom (blue). **I**. Boxplots showing relative exhaustion scores of CTL cells in different cancer types, with background color separating different studies.

**Figure S7.**
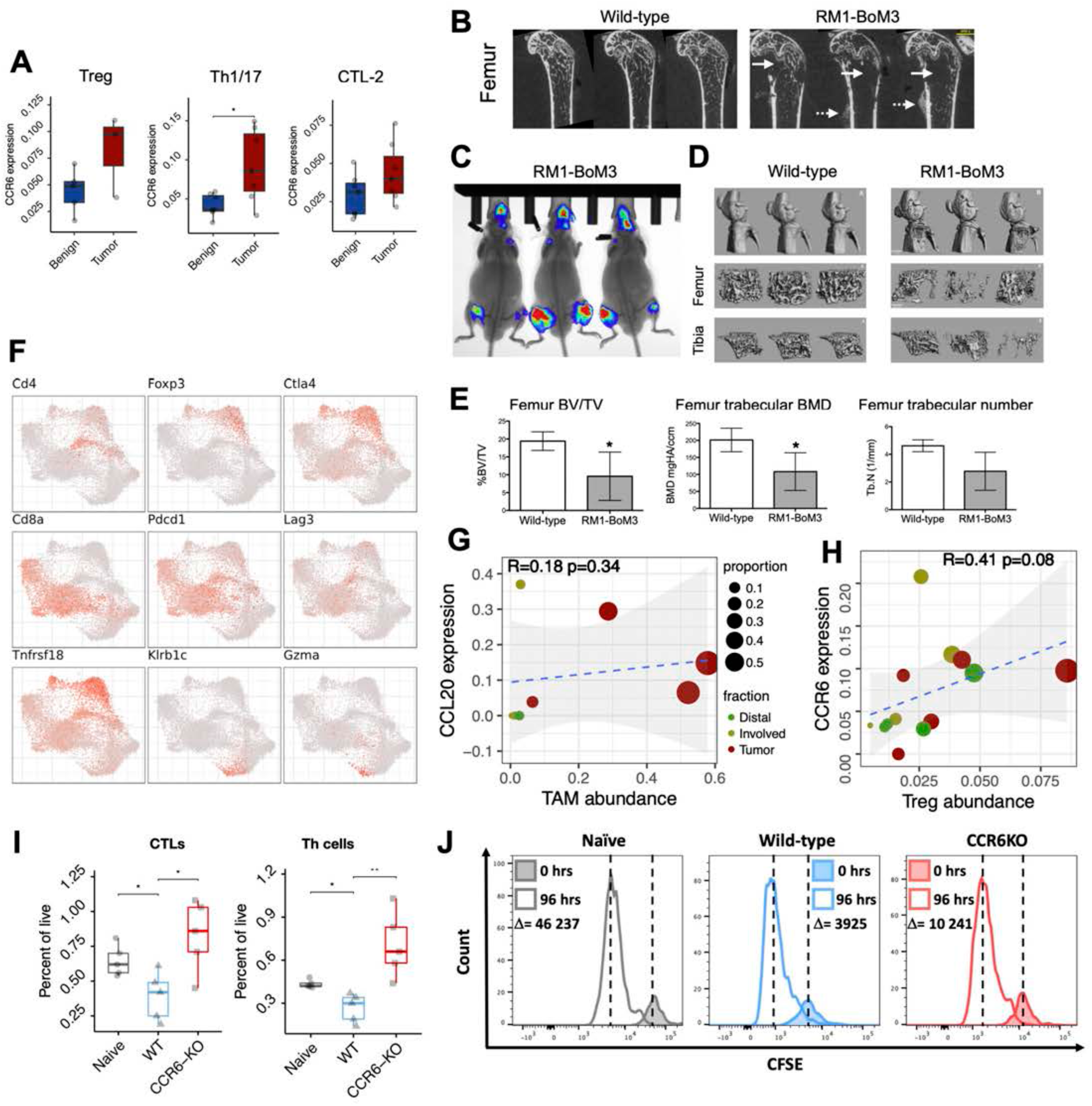
Disruption of the CCL20-CCR6 signaling axis relieves T cell exhaustion and improves survival. **A**. CCR6 expression levels in Treg and TH cells. **B**. Radiography imaging of femurs from wild-type mice without bone metastases and wild-type mice with RM1-BoM3 cells 3 weeks post intracardiac injection representing osteolytic lesions (solid arrows) and osteoblastic lesions (dotted arrow). **C**. Illustration of a third-generation syngeneic bone tropic prostate bone metastasis cell line expressing luciferase and Tdtomato allowing to monitor experimental bone metastases in live syngeneic mice by bioluminescence imaging. **D**. Micro-computed tomography of long bones from wild-type mice without bone metastases and wild-type mice injected with RM1-BoM3 cells 3 weeks post intracardiac injection (3 representative mice per group). **E**. Quantification of the osteolytic lesions induced by the RM1-BoM3 cells in femur of mice (n=4 to 5 mice per group)3 per group). Statistical analysis was performed using an unpaired Student’s t-test (*p<0.05). **F**. Selected marker gene expression in T cells of bone metastatic model. **G**. Similar to Figure 71, the plot illustrates correlation between the TAM abundance (assessed as a proportion of all myeloid cells in the sample) and CCL20 expression in TAM. **H**. Scatter plot illustrates correlation between Treg abundance and the CCR6 expression in each respective population (y axis). x axis assessed as a proportion of all T cells in the sample. I. Abundance of CTL and Th populations is shown as a percentage of all live cells (y axis) for Naive mice, and mice with bone marrow metastasis: WT and CCR6-KO. J. Shifts in the CFSE dye intensity between 0 and 96 hours is shown for the CD8+ T cell populations sorted from Naive, and metastasis-bearing WT and CCR6-KO mice. Vertical lines denotes MFI for individual samples.

